# Highly precise base editing with CC context-specificity using engineered human APOBEC3G-nCas9 fusions

**DOI:** 10.1101/658351

**Authors:** Zhiquan Liu, Siyu Chen, Huanhuan Shan, Mao Chen, Yuning Song, Liangxue Lai, Zhanjun Li

**Author notes:** To whom correspondence should be addressed; (Zhanjun Li); (Liangxue Lai). These authors contributed equally to this work (Zhiquan Liu, Siyu Chen, Huanhuan Shan).

## Abstract

Cytidine base editors, composed of a cytidine deaminase fused to Cas9 nickase, enable efficient C-to-T conversion in various organisms. However, current base editors can induce unwanted bystander C-to-T conversions when more than one C is present in the activity window of cytidine deaminase, which negatively affects the precision. Here, we develop a new base editor with CC context-specificity using rationally engineered human APOBEC3G, thus significantly reduce unwanted bystander activities. In addition, efficient C-to-T conversion that can further recognize relaxed NG PAMs is achieved by combining an engineered SpCas9-NG variant. These novel base editors with improved precision and targeting scope will expand the toolset for precise gene modification in organisms.

## Background

In contrast to conventional gene-editing nucleases, cytidine base editor (CBE) can achieve targeted C-to-T conversions without requiring DNA double-strand breaks (DSBs) or a donor template, and it induce lower levels of unwanted insertion/deletion mutations (indels), represent significant advances in precise genome manipulation^1^. However, the most common used base editor 3 (BE3), consists of rat APOBEC1 (rA1) fused with a *Streptococcus pyogenes* Cas9 (SpCas9) nickase and uracil glycosylase inhibitor (UGI), can induce unwanted bystander C-to-T conversions when more than one C is present in the enzyme’s activity window^2, 3^. It can negatively affects the precision of targeted base editing, thus not ideal for precise disease modelling and gene therapy when accurate single C substitution is required. To overcome this limitation, To overcome this limitation, optimized rA1 with mutant deaminase domains (YE base editors) or shortened linker has been used to effectively narrow the width of the editing window in human cells^4, 5^. Moreover, an engineered human APOBEC3A (eA3A) domain with TC context-specificity has been reported to efficiently reduce bystander mutations, and it has been proven superior to conventional base editors with narrowed window^6^. Context-dependent base editors represent an important advance that offers more precise base editing, while the application of eA3A-BEs was restricted by the presence of TCR (A/G) motifs^6^. Here, a new base editor was developed for drastically reducing bystander mutations using an engineered human APOBEC3G (hA3G) C-terminal catalytic domain, which preferentially deaminates cytidines in specific motifs according to a CC<u>C</u>≥C<u>C</u>C≥C<u>C</u> hierarchy firstly (where the preferentially deaminated C is underlined). Moreover, the further engineered variants, A3G-D316R/D317R and A3G-NG, could be used to induce more efficient base editing in CC motif and expand genome-targeting scope with NG PAMs.

## Results and Discussion

Previous study showed hA3G preferentially deaminates cytidines in C<u>C</u> and CC<u>C</u> motif *in vitro*^7^, thus it has the potential to develop as a CC context-dependent base editor. In addition, hA3G has a C-terminal catalytic domain and a second pseudocatalytic domain at N-terminal that retains the same tertiary fold, but are not catalytically active^7^. In this study, we replaced rA1 with the C-terminal catalytic domain of hA3G (hA3G-CTD) in BE4max^8^, current optimal architecture of CBE, to create A3G-BE4max (Fig. 1a). We first compared the ability of BE4max and A3G-BE4max to edit three previously reported target sites with multiple Cs in human cells (*EMX1, FANCF* and Site A) by co-transfecting with the respective single guide RNAs (sgRNAs) into 293T cells^4^ (Fig S1a). Base editing frequencies were evaluated from Sanger sequence chromatograms of each blastocyst using EditR, a robust and inexpensive base editing quantification software^9^. Genomic DNA analysis indicated the A3G-BE4max exhibited similar efficiency comparable to that of BE4max at three sites, meanwhile showed obvious preference in C<u>C</u> (*EMX1*) and C<u>CC</u> (*FANCF* and Site A) contexts (Fig S1b-d).

**Figure 1.**
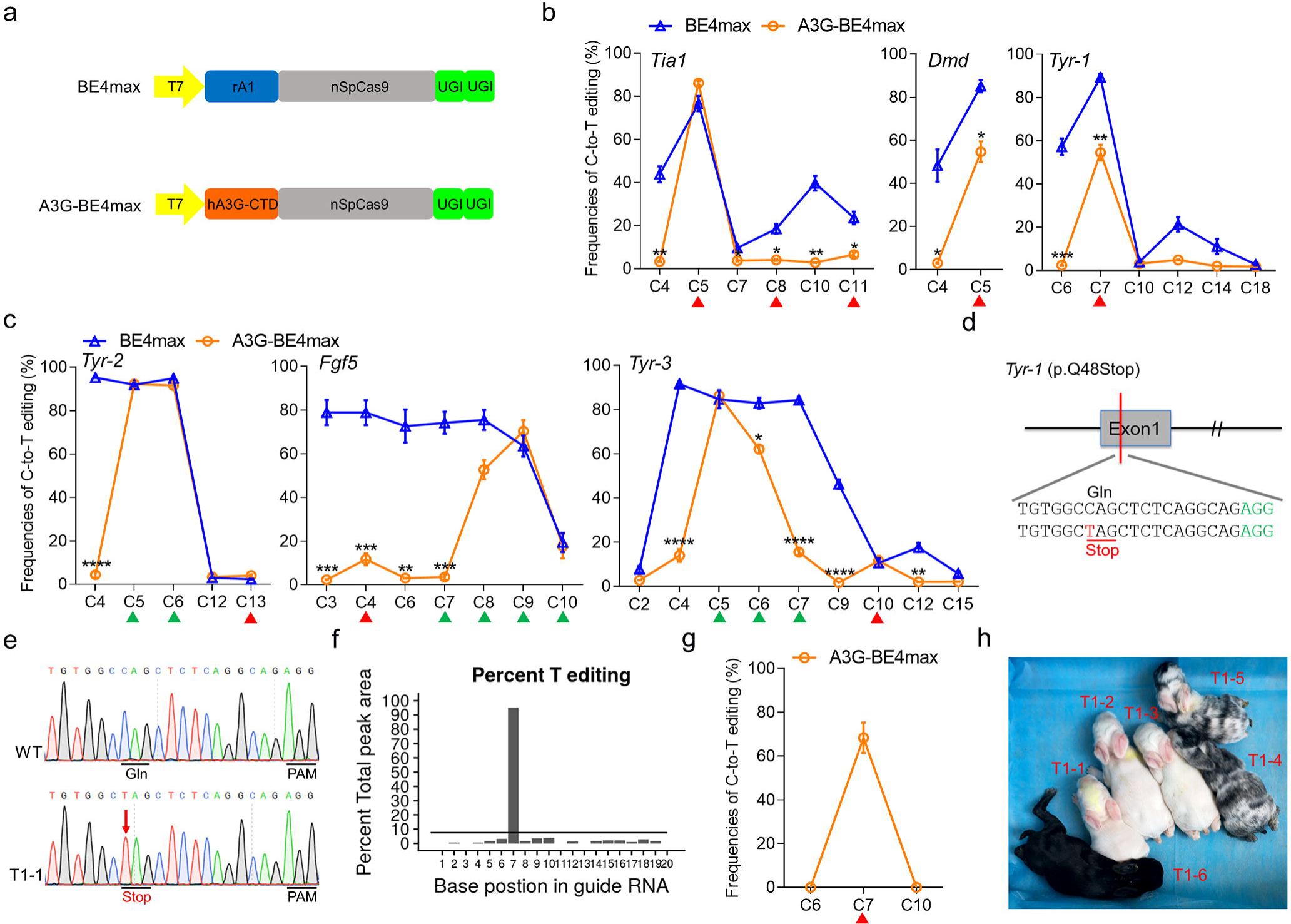
A3G-BE4max can induce efficient C-to-T conversions with minimized bystander activity. **a** Schematic representation of BE4max and A3G-BE4max architecture. **b, c** Frequencies of single C-to-T conversions using BE4max and A3G-BE4max at six sites with CC (**b**) or CCC (**c**) motif in rabbit embryos. C<u>C</u> context (red triangle) and C<u>CC</u> context (green triangle). Target Cs are counting the PAM as positions 21-23. **d** Target sequence at the *Tyr-1* (p.Q48stop) locus. PAM region (green) and target mutation (red). **e** Representative sequencing chromatograms from a WT, and mutant rabbit (T1-1). The red arrow indicates the substituted nucleotide. The relevant codon identities at the target site are presented under the DNA sequence. **f** The predicted editing bar plot based on Sanger sequencing chromatograms from T1-1 by EditR. **g** Single C-to-T editing frequencies of F0 rabbits at *Tyr-1* using A3G-BE4max. **h** Photograph of all six F0 rabbits at 1 week.

To further evaluate the precision of this system *in vivo*, six target sites including CC and CCC contexts were selected to test in rabbit embryos (Table S1). The control group, BE4max, showed a large editing window mainly from C3 to C9 (∼7 nt) and even induced the widest range of mutations spanning from C2 to C15 at *Tyr-3* without obvious context-specificity (Fig. 1b, 1c and : Fig S2 and S4). In contrast, A3G-BE4max exhibited ideal efficiency comparable to that of BE4max at all six sites (average editing frequencies: 54.52-92.24% vs. 76.55-95.20%), but significantly reduced bystander activities in non-C<u>C</u> or C<u>CC</u> contexts with a similar ∼7 nt editing window (mainly from C4 to C10) (Fig. 1b, 1c). Moreover, the A3G-BE4max exclusively edit the second C when the CC dinucleotide presents in the editing window, while with reduced efficiency at two (54.68 ± 10.70% vs. 85.12 ± 6.34%, *p* < 0.05 at *Dmd*; 54.52 ± 8.82% vs. 89.28 ± 4.19%, *p* < 0.01) of three tested sites compared with BE4max (Fig. 1b). With the high precision of A3G-BE4max, targeted C-to-T conversions can be induced at target C without generating bystander mutations at *Tia1*, thus precisely mimic the p.P362L missense mutation of human Amyotrophic Lateral Sclerosis (ALS)^10^ (Fig. 1b and S3). In addition, at three tested sites with multiple Cs, A3G-BE4max induced high editing frequencies in C<u>CC</u> (*Tyr-2* and *Tyr-3*) or CC<u>C</u> (*Fgf5*) contexts, meanwhile notably decreased bystander activities at the first cytidine (Fig. 1c). Overall, these results demonstrated that engineered hA3G (hA3G-CTD) can efficiently possess its cytidine deaminase sequence preference of C<u>C</u> in the context of a base editor fusion (A3G-BE4max).

Next, a single C-to-T conversion was designed at *Tyr-1* (p.Q48stop) to mimic human oculocutaneous albinism type 1 (OCA1) in Founder (F0) rabbits^11^ (Fig. 1d). The result of T-A cloning showed that five of six pups (83%) were mutants with editing frequencies from 40% to 100% (Fig. 1e, 1f and Fig S5, Table S2). Notably, targeted base editing at C7 was successfully induced in all of five mutants without any bystander mutations, enabling it to induce highly precise p.Q48Stop mutation of OCA1 (Fig. 1g and Fig S5). Moreover, three homozygous mutants (T1-1, T1-2 and T1-3) exhibited a complete albino phenotype, while the chimeric mutants (T1-4 and T1-5) exhibited mosaic distribution of black and white skin and hair, which is consistent with their mutant genotype (Fig. 1h and : Fig S6).

Additionally, considering that the variable efficiency at target sites with CC motif by A3G-BE4max (Fig. 1b), and the previous findings that mutations of residue D316R/D317R, D131E or N244G may modulate catalytic activity and sequence preference of hA3G^12, 13^, we introduced these mutations into A3G-BE4max to create A3G-D316R/D317R, A3G-D316E and A3G-N244G. Two target sites (*Fgf5* and *Tyr-3*) that harbor CC and CCC motif simultaneously were selected to proof the concept (Fig. S7). Remarkably, A3G-D316R/D317R greatly increased the editing frequencies of C<u>C</u> context at C4 of *Fgf5* and C10 at *Tyr-3*, while retaining lower bystander activities in the non-CC or CCC motif (Fig. S7 and S8). Furthermore, the similar maximal editing efficiencies of target Cs as BE4max were obtained using A3G-D316R/D317R at both *Dmd* and *Tyr-1*, despite a slightly increased bystander mutations were observed (Fig. 2a, 2b and Fig. S8). Moreover, A3G-D316E further improved the precision to single base (mainly C9 at *Fgf5* and C5 at *Tyr-3*) and drastically decreased catalytic activities were observed in A3G-N244G, consistent with previous reports *in vitro*^13^ (Fig. S7 and S8).

**Figure 2.**
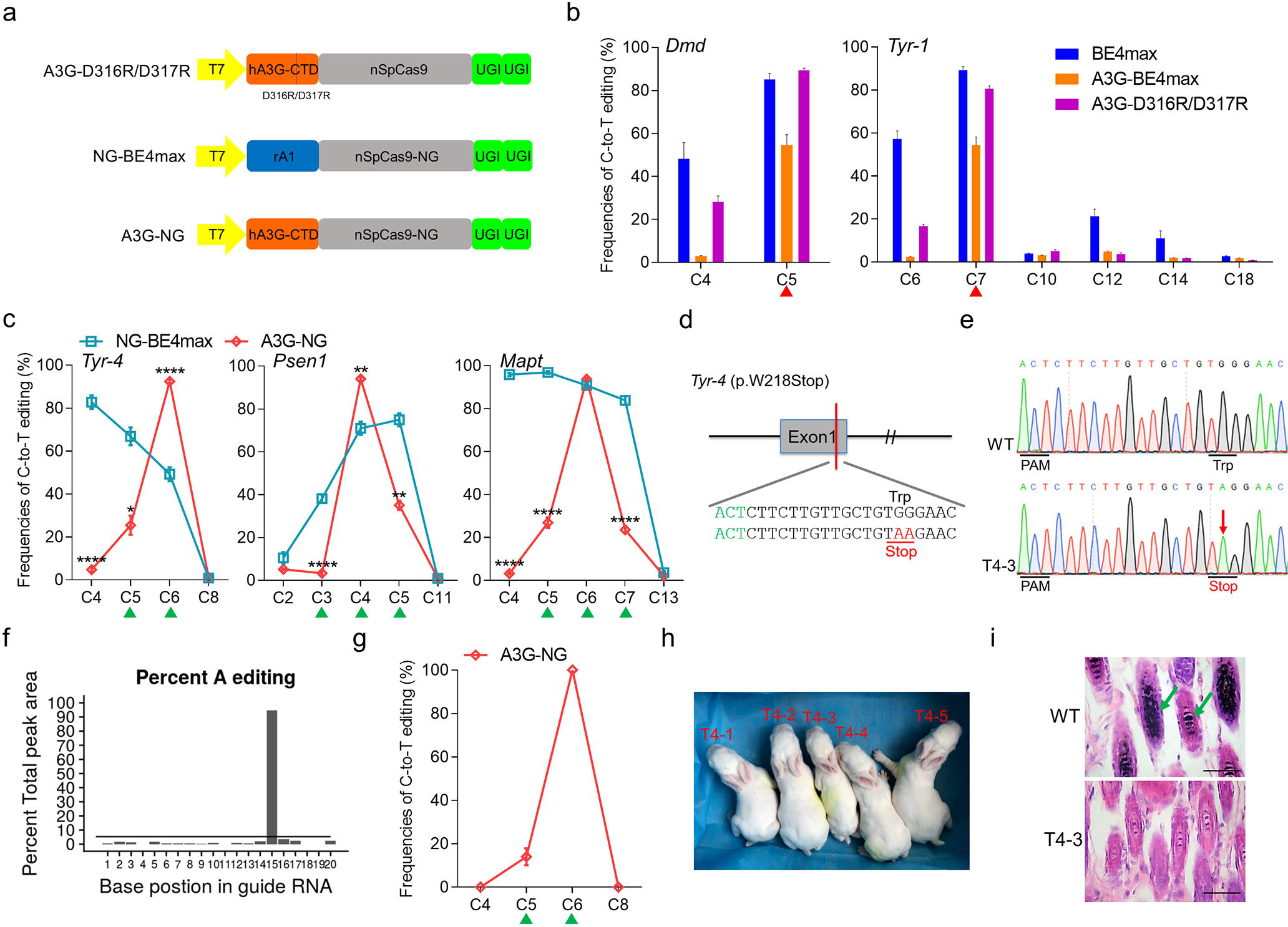
Enhanced efficiency and expanded targeting scope by optimized hA3G-nCas9 fusions. **a** Schematic representation of A3G-D316R/D317R, NG-BE4max and A3G-NG architecture. **b** Summary of C-to-T editing frequencies induced by three systems on each cytosine at *Dmd* and *Tyr-1*. C<u>C</u> context (red triangle). **c** Comparison of the editing frequencies of single C-to-T conversions between NG-BE4max and A3G-NG at three sites with NG PAMs in rabbit embryos. C<u>CC</u> context (green triangle). **d** Target sequence at the *Tyr-4* (p.W218Stop) locus. PAM region (green) and target mutation (red). **e** Representative sequencing chromatograms from a WT, and mutant rabbit (T4-3). The red arrow indicates the substituted nucleotide. The relevant codon identities at the target site are presented under the DNA sequence. **f** The predicted editing bar plot based on Sanger sequencing chromatograms from T4-3 by EditR. **g** Single C-to-T editing frequencies of F0 rabbits at *Tyr-4* (p.W218Stop) using A3G-NG. **h** Photograph of all five F0 rabbits at 1 week. **i** H&E staining of dorsal skin from WT and T4-3 rabbits. The green arrows highlight the melanin in the basal layer of the epidermis of WT rabbit. Scale bars: 50 μm.

In addition, the NGG PAM requirement of SpCas9 substantially limits the target sites suitable for hA3G-nCas9 fusions. Therefore, we explore the feasibility of SpCas9-NG system with currently the most relaxed NGN PAMs to expand the genome-targeting scope^14^ (Fig. 2a). Three target sites with NGT or NGA PAM were selected (Table S1). Notably, A3G-NG exhibited comparable high efficiency to that of NG-BE4max at all three sites of target Cs while significantly reduced bystander activities in non-C<u>CC</u> contexts, thus substantially decreased unwanted bystander mutations (Fig. 2c and S9 and S10). In particular, with both high precision and expanded space of A3G-NG, accurate p.P301L mutation can be induced in *Mapt* gene to precisely mimic human classical missense mutation of Alzheimer’s disease (AD)^15^. It is extremely difficult for conventional BE4max that can induce wrong mutation (p.P301F) due to high frequencies of bystander C-to-T editing (Fig. S10). Furthermore, the optimized A3G-NG was used to generate F0 rabbits that carry the *Tyr-4* (p.W218Stop) mutation in order to mimic human OCA1 (Fig. 2d). Five pups were obtained and all of them (100%) were homozygous with desired nonsense mutation, consistent with the high efficiency in embryos (Fig. 2e, 2f and Fig. S11, Table S2). Strikingly, no bystander mutations were observed in all rabbits (Fig. 2g and Fig. S10). All five pups (100%) showed a systemic albino phenotype (Fig. 2h). Moreover, histological H&E staining revealed the absence of melanin in the skin of representative T4-3, but not in the WT rabbit (Fig. 2i). These results suggested that it may be possible to further modify A3G-BEmax using a set of mutations in deaminase or Cas9 to tune precision and targeting scope.

## Conclusions

In summary, we develop a series of hA3G-nCas9 fusions that can induce efficient base editing with minimized bystander activity in CC and CCC motifs. The A3G-BE4max can function as a generic version of the CC context-dependent base editor, the A3G-D316R/D317R shows higher editing frequencies in CC dinucleotides, and the engineered A3G-NG further expands genome-targeting scope. Thus, these hA3G-nCas9 base editors improve the precision and expand the scope of the currently used rA1-BEs system.

## Methods

### Ethics statement

New Zealand white and Lianshan black rabbits were obtained from the Laboratory Animal Center of Jilin University (Changchun, China). All animal studies were conducted according to experimental practices and standards approved by the Animal Welfare and Research Ethics Committee at Jilin University.

### Plasmid construction

BE4max were obtained from Addgene (#112093). The DNA fragement of hA3G-CTD was synthesized and cloned into BE4max by Genscript Biotech (Nanjing) to create A3G-BE4max. The D316R/D317R, D316E or N244G mutations were induced into hA3G-CTD to produce A3G-D316R/D317R, A3G-D316E and A3G-N244G. Seven mutations (R1335A/L1111R/D1135V/G1218R/E1219F/A1322R/T1337R) of SpCas9 were introduced into BE4max and A3G-BE4max to create NG-BE4max and A3G-NG. Plasmid site-directed mutagenesis was performed using the Fast Site-Directed Mutagenesis Kit (TIANGEN, Beijing). All the site-directed mutation primers are listed in Table S3. The amino acid sequences of plasmids are listed in the Supplementary sequence.

### Cell culture and transfection

Human kidney epithelial cell line (HEK293T) were cultured in Dulbecco’s modified Eagle’s medium (DMEM) supplemented with 10% fetal bovine serum (HyClone), and incubated at 37°C in an atmosphere of 5% CO2. The cells were seeded into 6-well plates and transfected using LipofectamineTM 3000 Reagent (Thermo Fisher Scientific) according to the manufacturer’s instructions. After 72 hours, the cells were collected and used for genotyping. All primers used for genotyping are listed in Table S5.

### mRNA and gRNA preparation

All plasmids were linearized with NotI and transcribed in vitro using the HiScribe(tm) T7 ARCA mRNA kit (NEB). mRNA was purified using the RNeasy Mini Kit (Qiagen) according to the manufacturer’s protocol. sgRNA oligos were annealed into pUC57-sgRNA expression vectors containing a T7 promoter. The sgRNAs were then amplified and transcribed *in vitro* using the MAXIscript T7 kit (Ambion) and purified using the miRNeasy Mini Kit (Qiagen) according to the manufacturer’s protocol. The sgRNA oligo sequences used in this study are listed in Table S4.

### Microinjection of rabbit zygotes

The protocol used for the microinjection of pronuclear-stage embryos has been described in detail in our previously published study^3^. Briefly, a mixture of mRNA (200 ng/ul) and sgRNA (50 ng/ul) was co-injected into the cytoplasm of pronuclear-stage zygotes. The injected embryos were transferred into EBSS medium for short-term culture at 38.5 °C, 5% carbon dioxide and 100% humidity. Then approximately 30–50 injected zygotes were transferred into the oviducts of recipient rabbit.

### Single-embryo PCR amplification and rabbit genotyping

Each group was injected with an average of approximately 10 embryos to test the base editing efficiency. The injected embryos were transferred into EBSS medium for culture at 38.5 °C, 5% carbon dioxide and 100% humidity. Then the injected embryos were collected at the blastocyst stage. Genomic DNA was extracted in embryo lysis buffer (1% NP40) at 56 °C for 60 minutes, then at 95 °C for 10 minutes in a BIO-RAD PCR Amplifier. Then the extracted products were amplified by PCR (95°C, 5min for pre-degeneration, 42 cycles of (95°C, 30s, 58°C, 30s, 72°C, 30s), 72°C, 5min for extension) and determined by Sanger sequencing. The Sanger sequencing result of each blastocyst was used to evaluate base editing frequencies by EditR^9^. The genomic DNA of newborn rabbits was extracted from ear clips and analysed by PCR genotyping, Sanger sequencing and T-A cloning. All primers used for genotyping are listed in Table S5.

### Haematoxylin and eosin (H&E) staining

The dorsal skin from WT and mutant rabbits was fixed in 4% paraformaldehyde for 48 hours, embedded in paraffin wax and then sectioned for slides. Slides were stained with H&E and viewed under a Nikon ts100 microscope.

### Statistical analysis

All data are expressed as mean ± SEM of at least three individual determinations for all experiments. Data were analyzed by student’s t-test via GraphPad prism software 6.0. The probability value that smaller than 0.05 (p < 0.05) was considered to be statistically significant. \**p* < 0.05, \*\**p* < 0.01, \*\*\**p* < 0.001, \*\*\*\**p* < 0.0001.

## List of abbreviations

(CBE): (cytidine base editor);
(DSBs): (double-strand breaks);
(indels): (insertion/deletion mutations);
(BE3): (base editor 3);
(rA1): (rat APOBEC1);
(SpCas9): (*Streptococcus pyogenes* Cas9);
(UGI): (uracil glycosylase inhibitor);
(eA3A): (engineered human APOBEC3A);
(hA3G): (human APOBEC3G);
(sgRNAs): (single guide RNAs);
(OCA1): (oculocutaneous albinism type 1);
(F0): (Founder);
(AD): (Alzheimer’s disease);
(H&E): Haematoxylin and eosin.

## Declarations

### Acknowledgements

The authors thank Peiran Hu and Nannan Li for assistance at the Embryo Engineering Center for critical technical assistance.

### Conflict of interests

The authors declare no competing interests.

### Funding

This study was financially supported by the National Key Research and Development Program of China Stem Cell and Translational Research (2017YFA0105101). The Program for Changjiang Scholars and Innovative Research Team in University (No.IRT_16R32). The Strategic Priority Research Program of the Chinese Academy of Sciences (XDA16030501, XDA16030503), Key Research & Development Program of Guangzhou Regenerative Medicine and Health Guangdong Laboratory (2018GZR110104004).

### Authors’ contributions

ZL, LL and ZL conceived and designed the experiments. ZL, SC and HS performed the experiments. ZL, HS, SC analysed the data. MC and YS contributed reagents/materials/analysis tools. ZL and ZL wrote the paper. All authors have read and approved the final manuscript.

### Data availability

The authors state that all data necessary for confirming the conclusions presented in this article are represented fully within the article, or can be provided by the authors upon request.

