## Supplementary materials for "Highly precise base editing with CC context-specificity using engineered human APOBEC3G-nCas9 fusions"

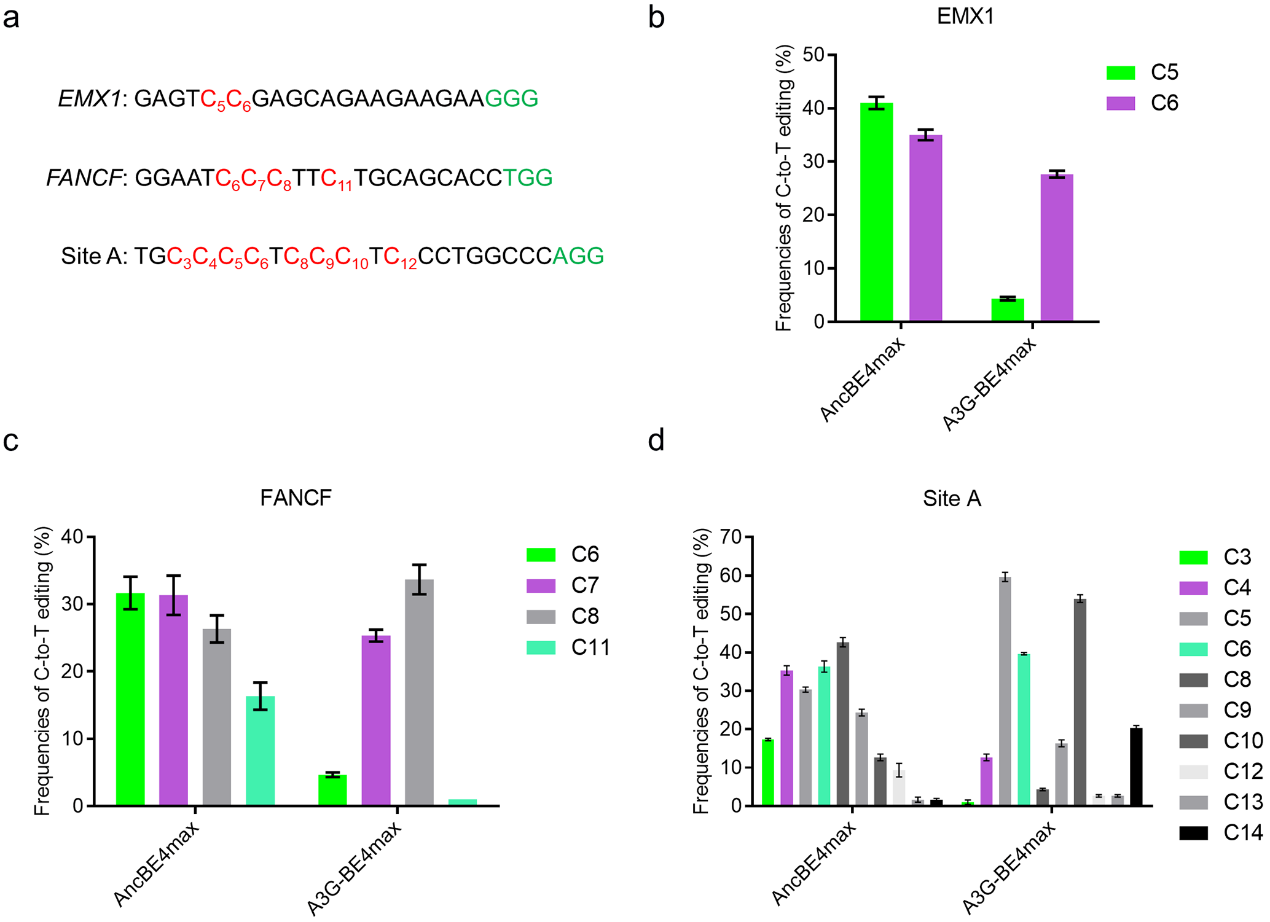

**Figure S1.** Comparison of C-to-T editing efficiency in HEK293T cells using BE4max and A3G-BE4max. **a** Protospacers and PAM (green) sequences of the genomic loci tested, with the target Cs shown in red. Cytosines are counted with the base distal to the PAM setting as position 1. **b-d** Summary of C-to-T editing frequencies induced by two systems on each cytosine at three sites. The editing frequencies were evaluated by EditR.

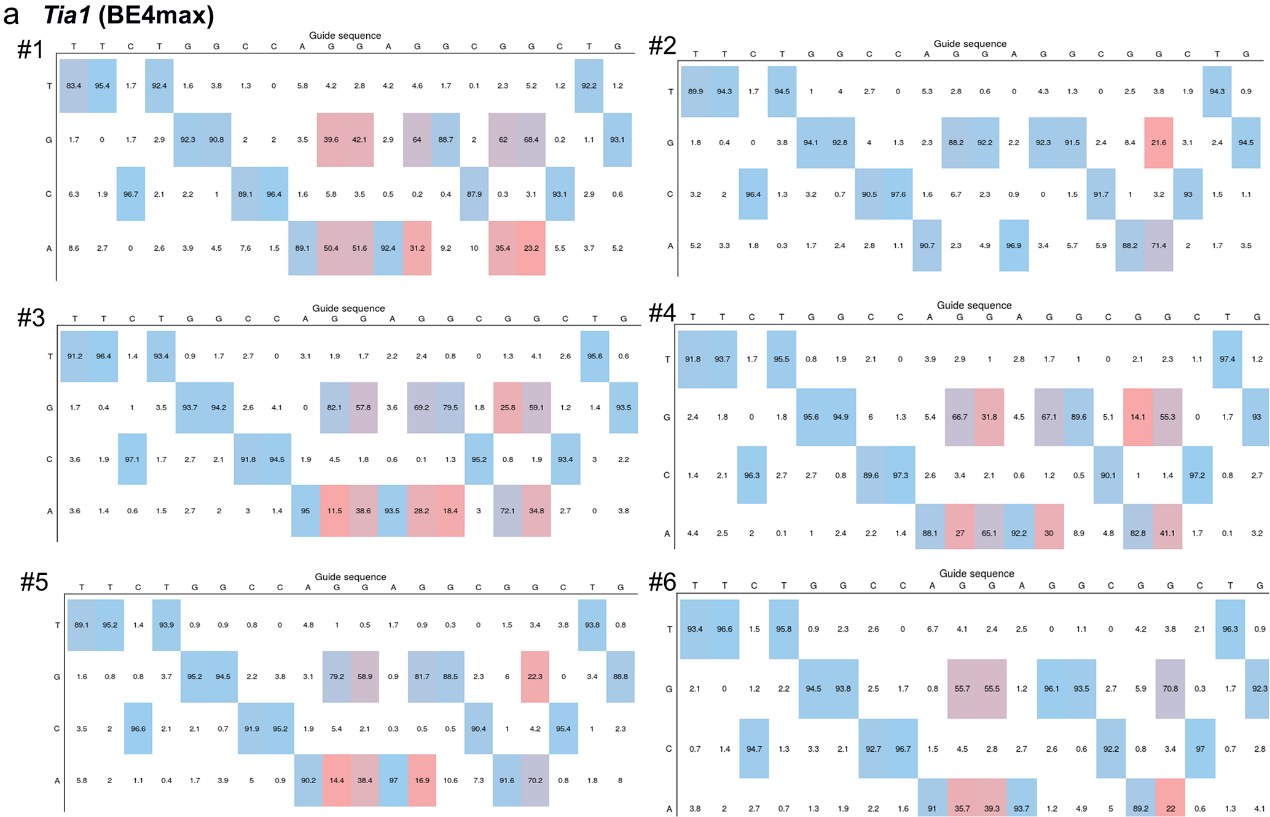

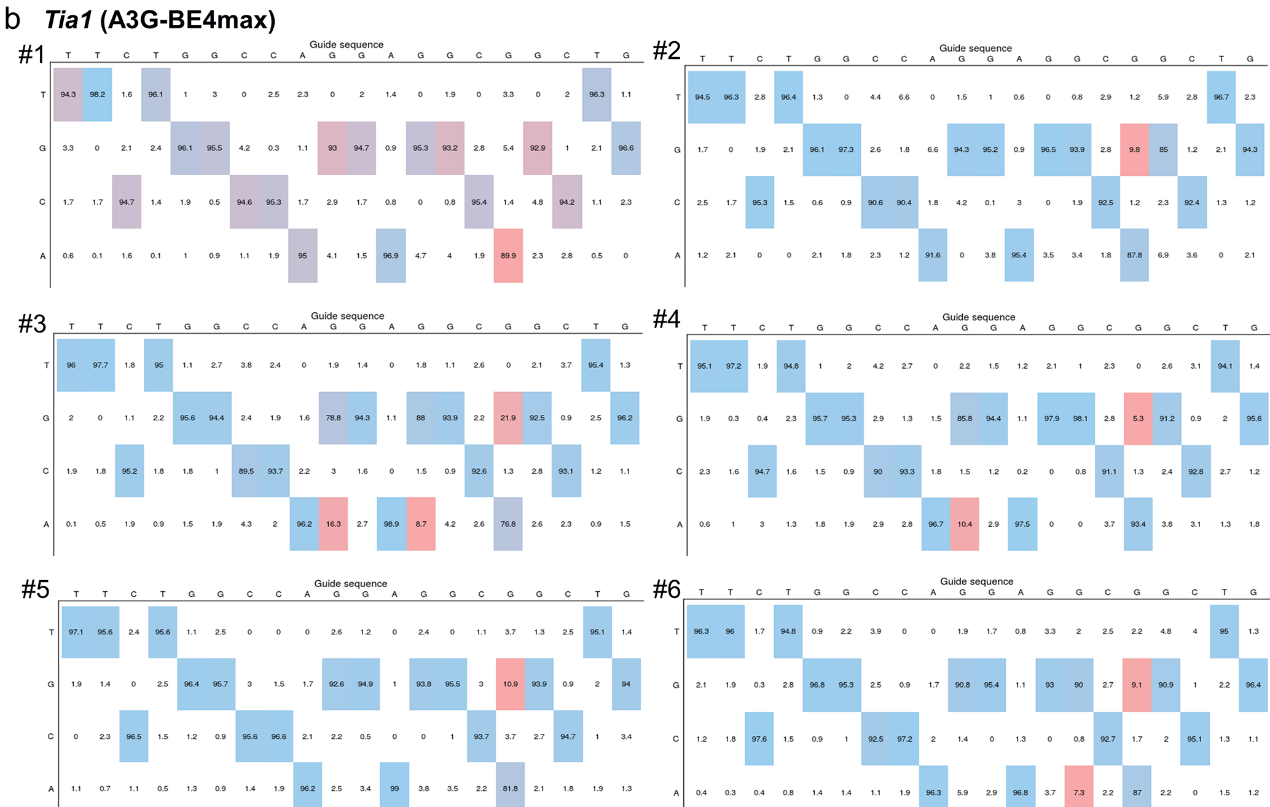

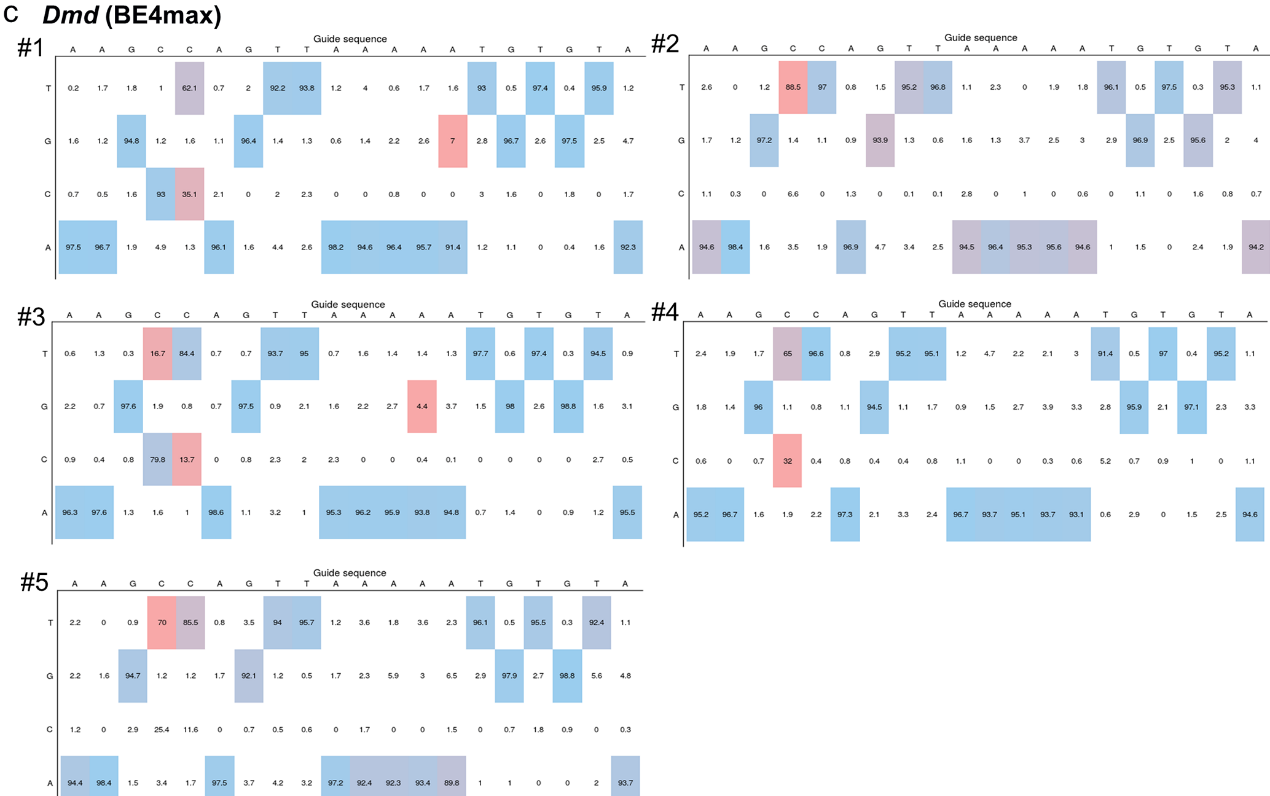

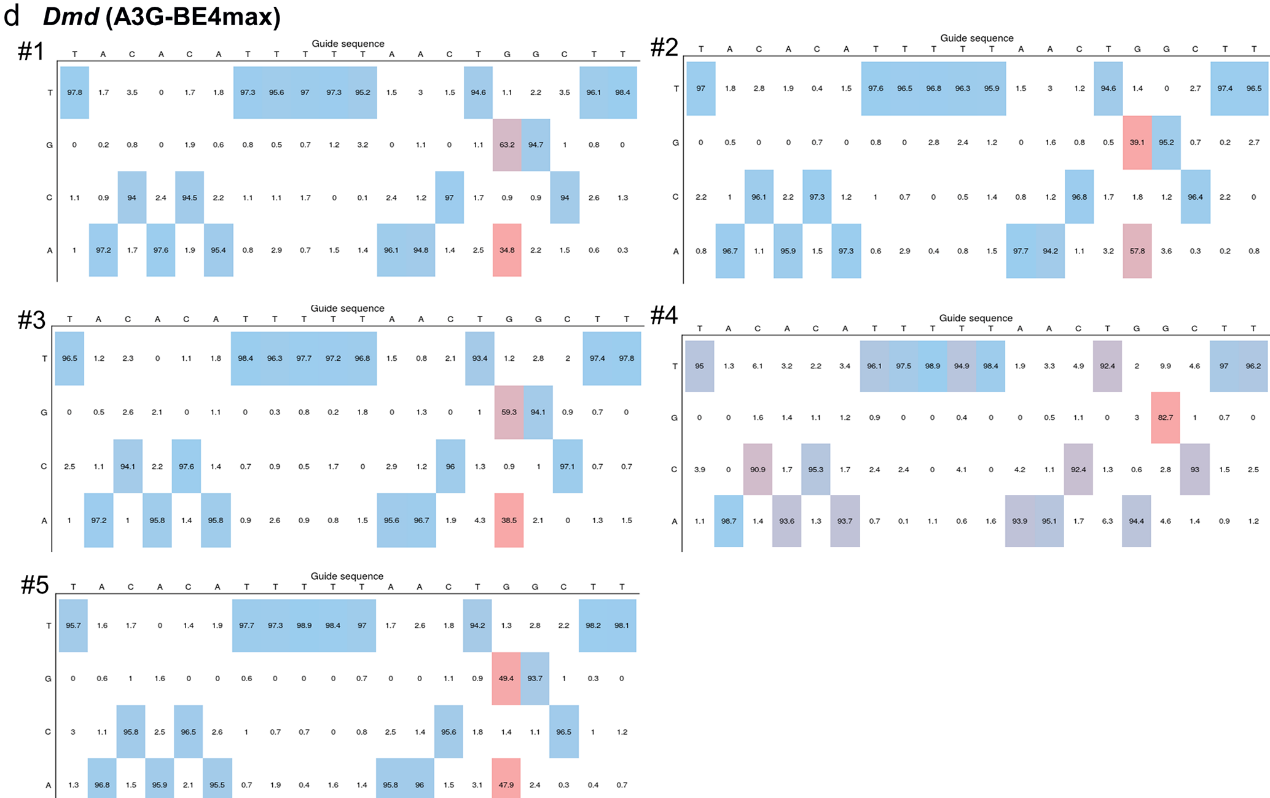

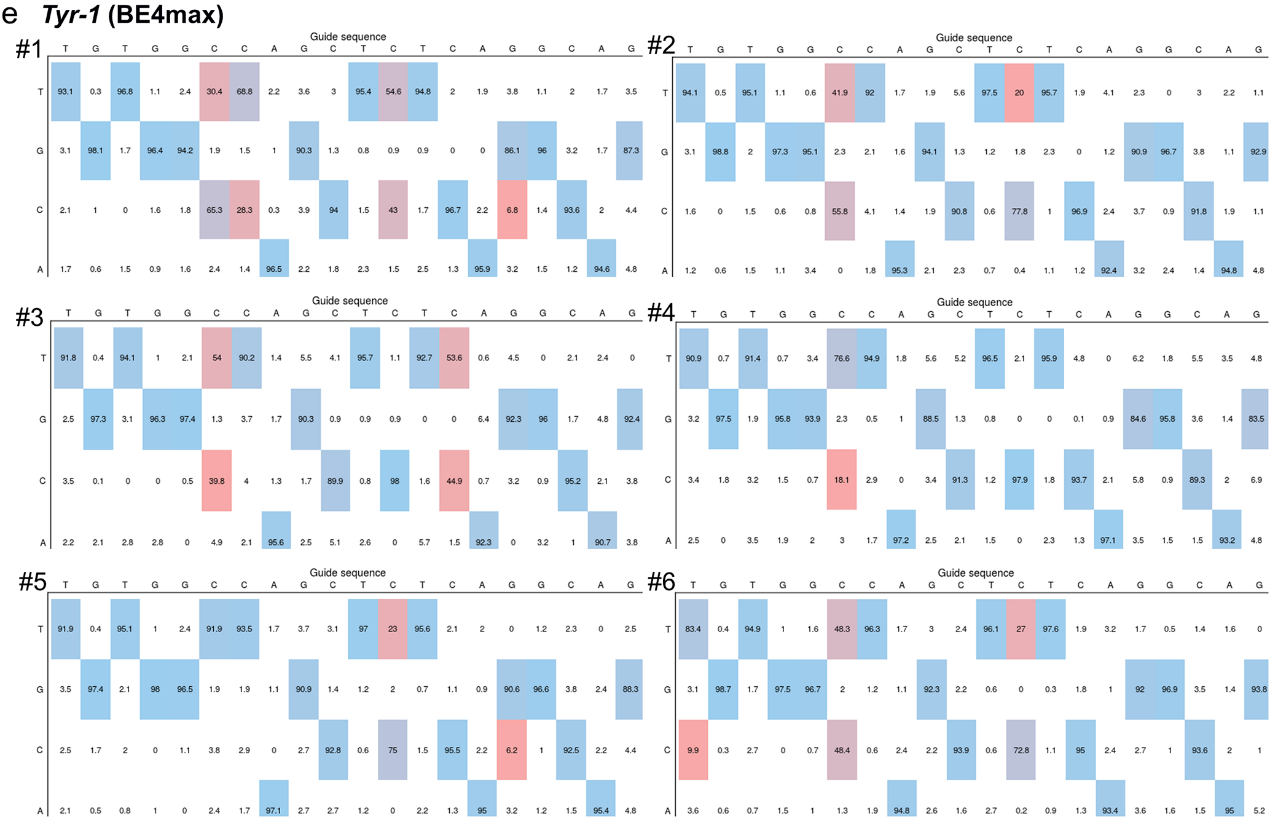

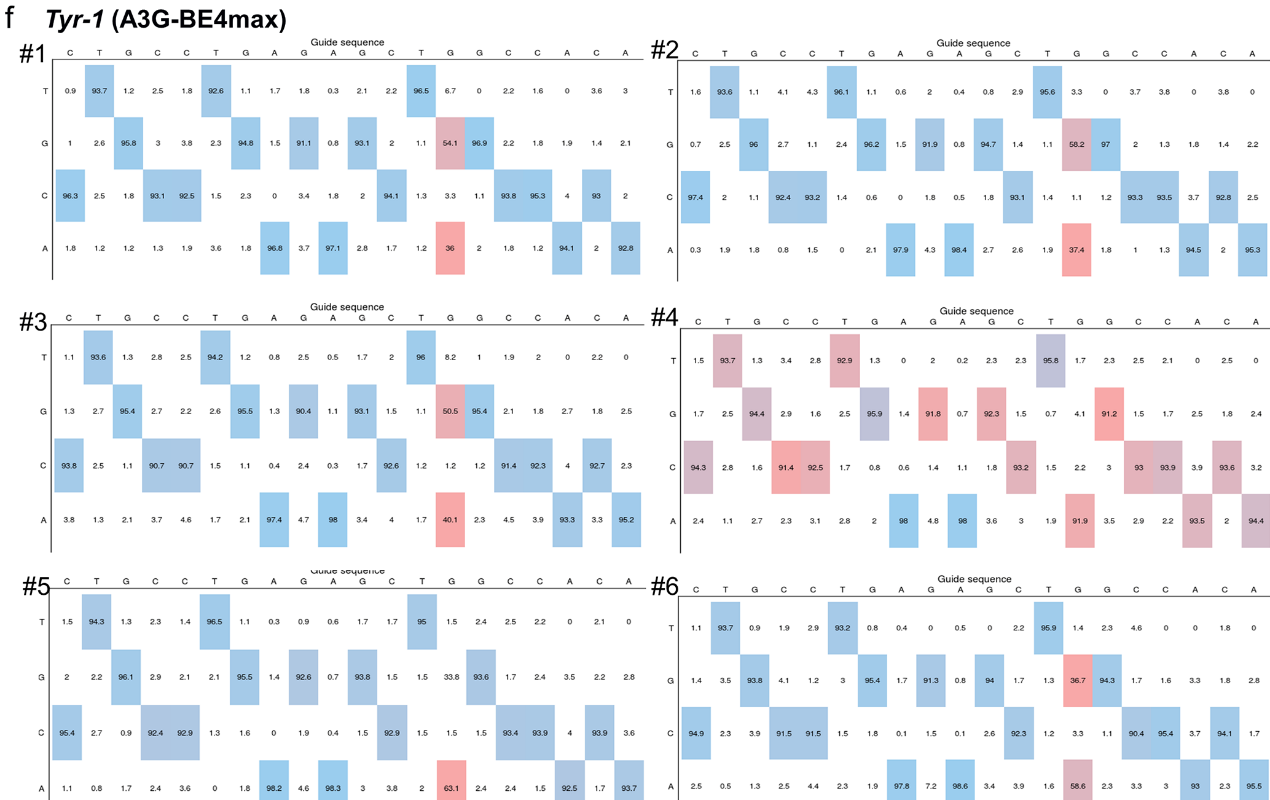

**Figure S2.** Base editing frequencies and product distribution at *Tia1* (**a**, **b**), *Dmd* (**c**, **d**) and *Tyr-1* (**e**, **f**) sites with CC context in rabbit blastocysts using BE4max and A3G-BE4max. The output editing tables in a colored heat map were produced by evaluating the Sanger sequencing results of each blastocyst by EditR.

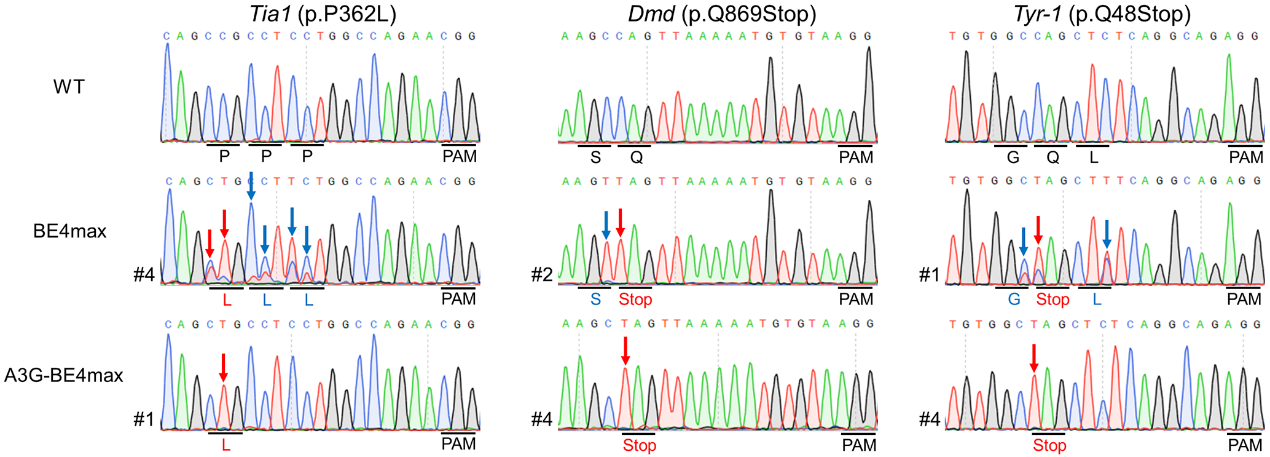

**Figure S3.** Representative sequencing chromatograms of edited rabbit blastocyst at three target sites using BE4max and A3G-BE4max systems. Targeted mutations (red arrows) and bystander mutations (blue arrows). The relevant codon identities at the target site are presented under the DNA sequence.

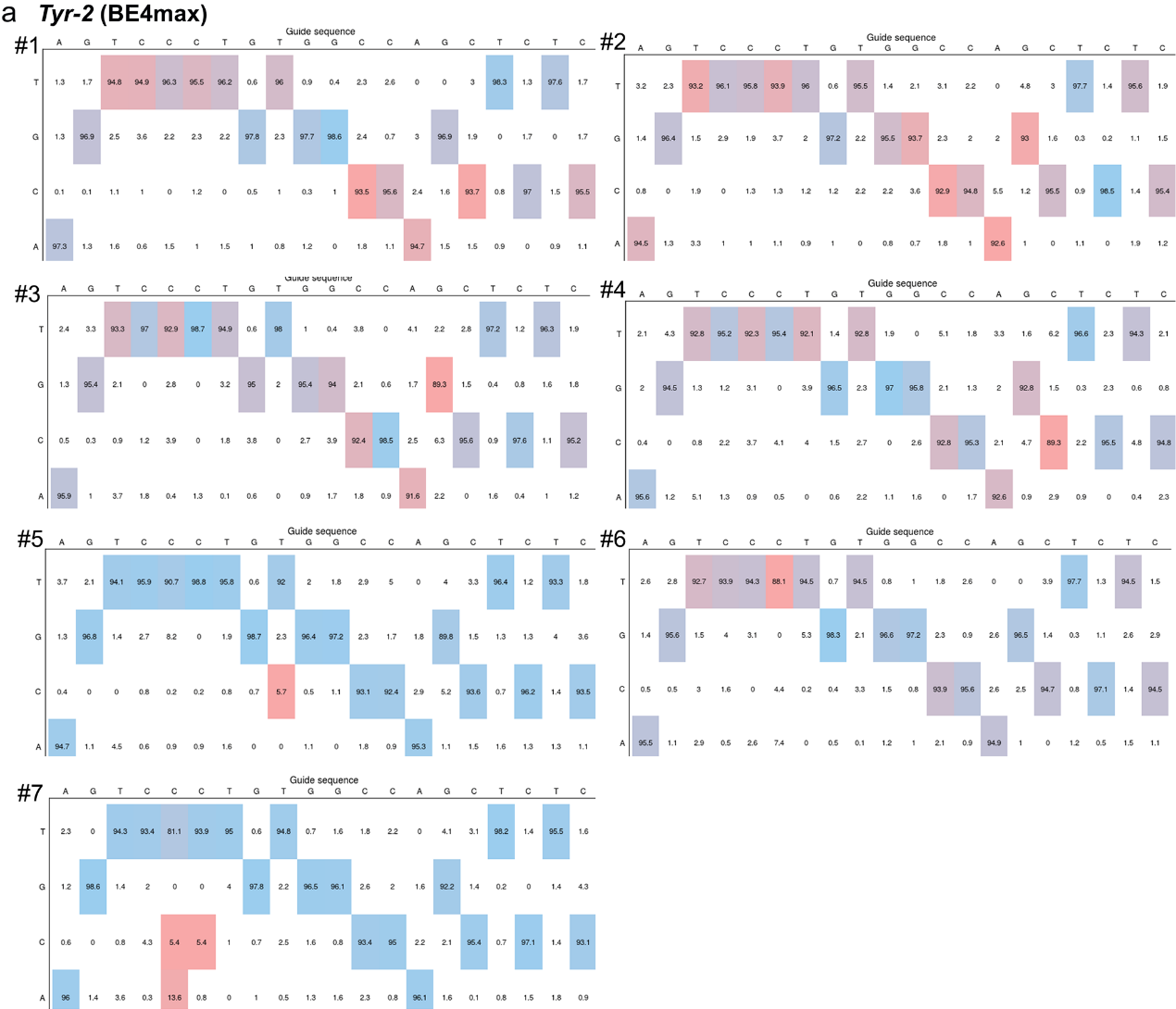

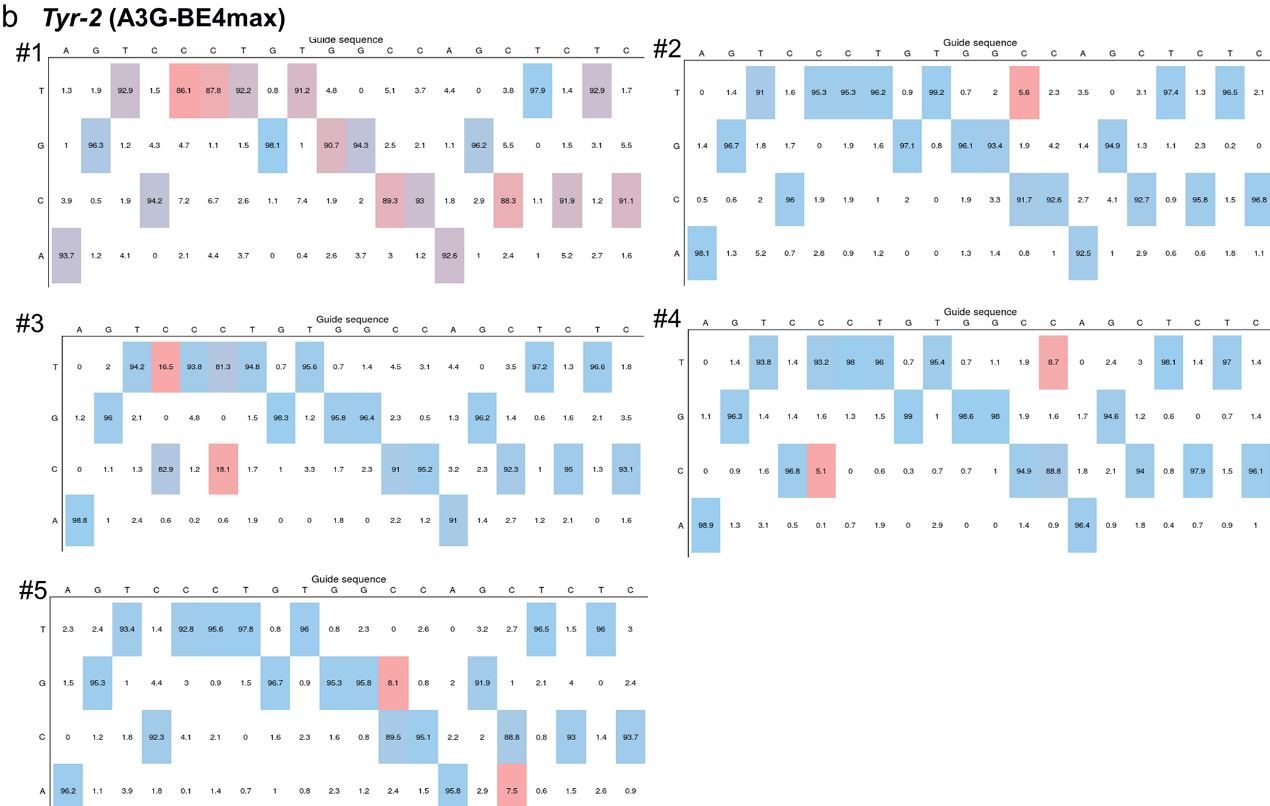

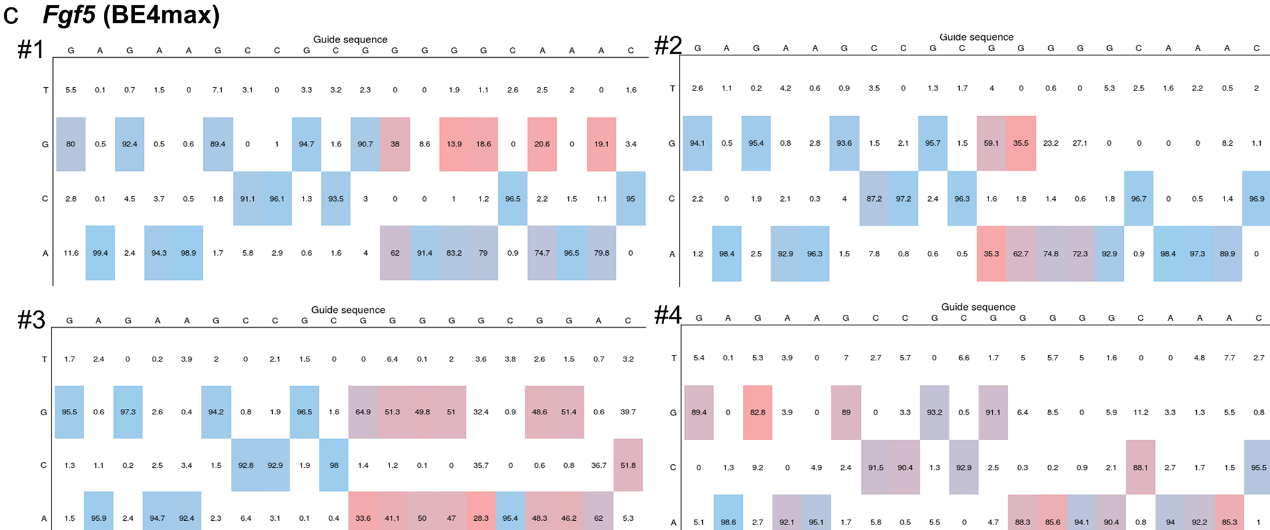

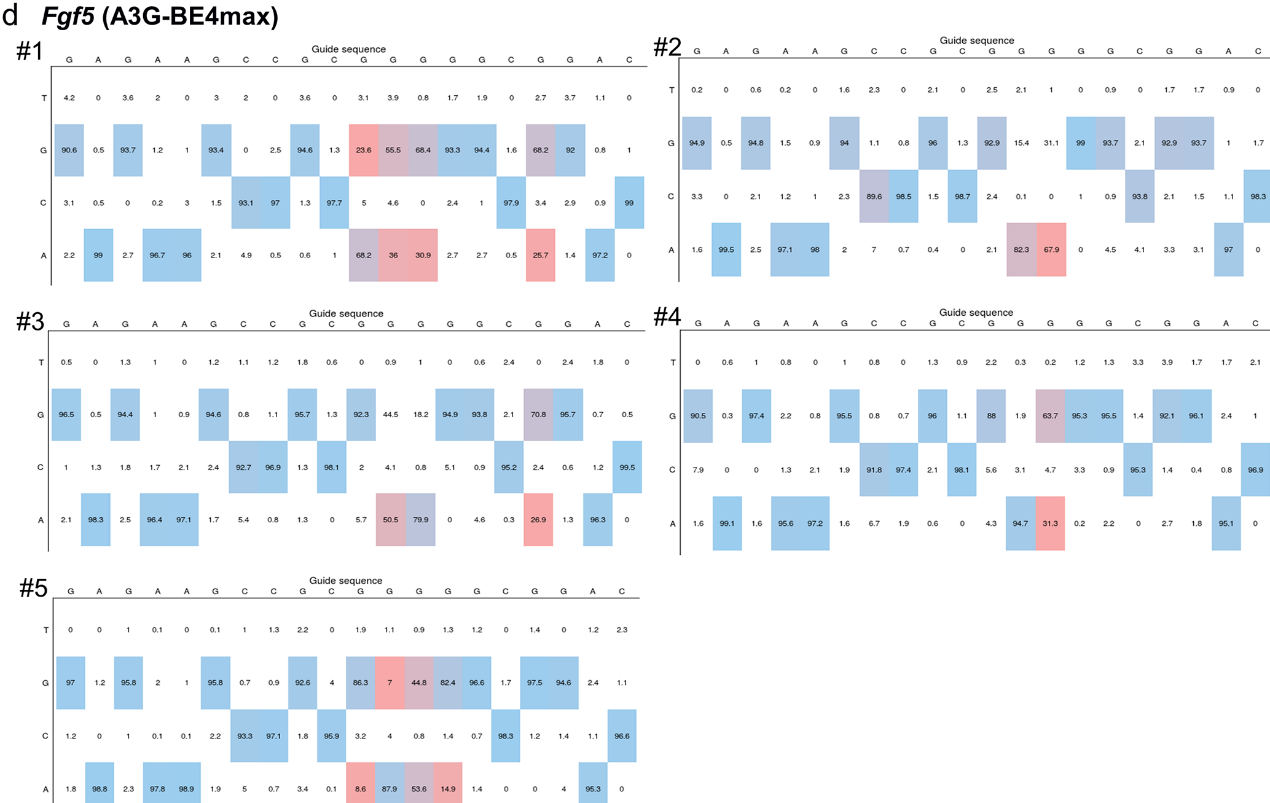

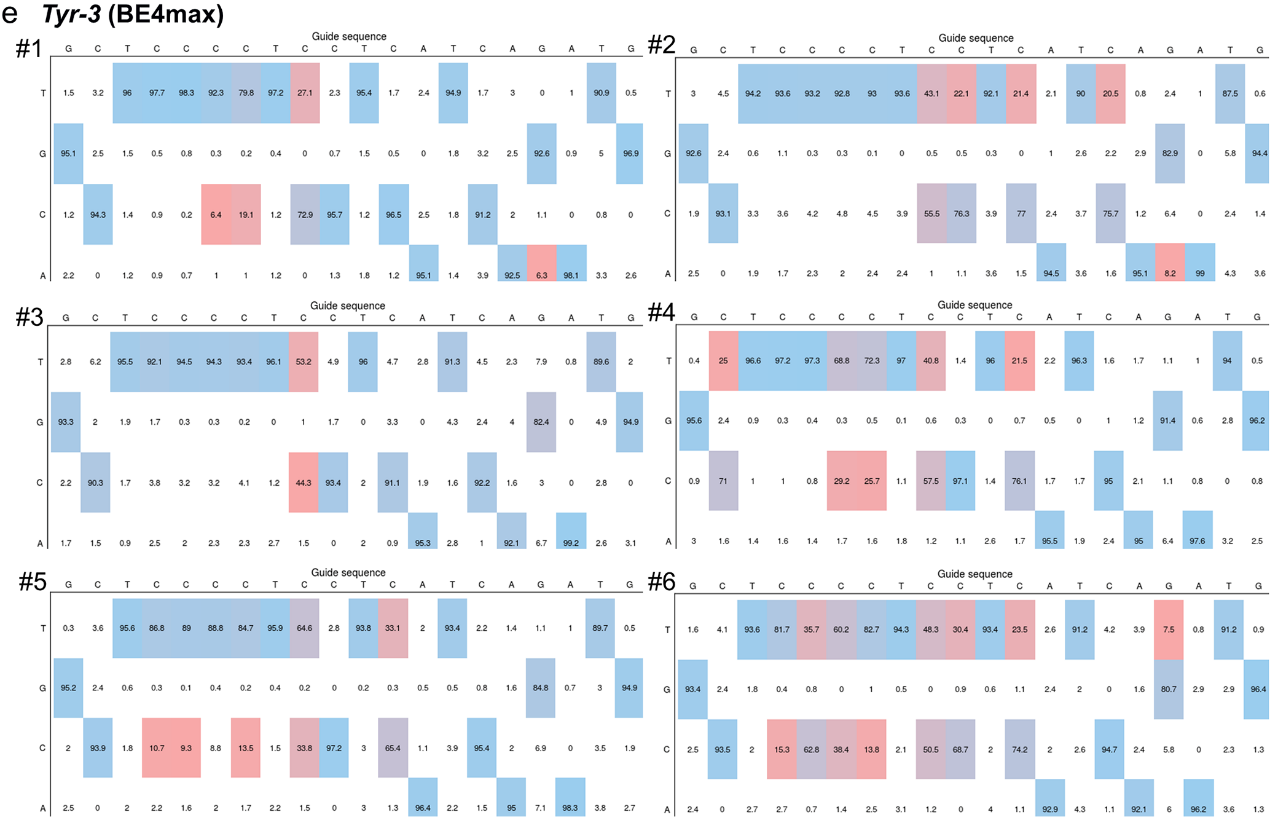

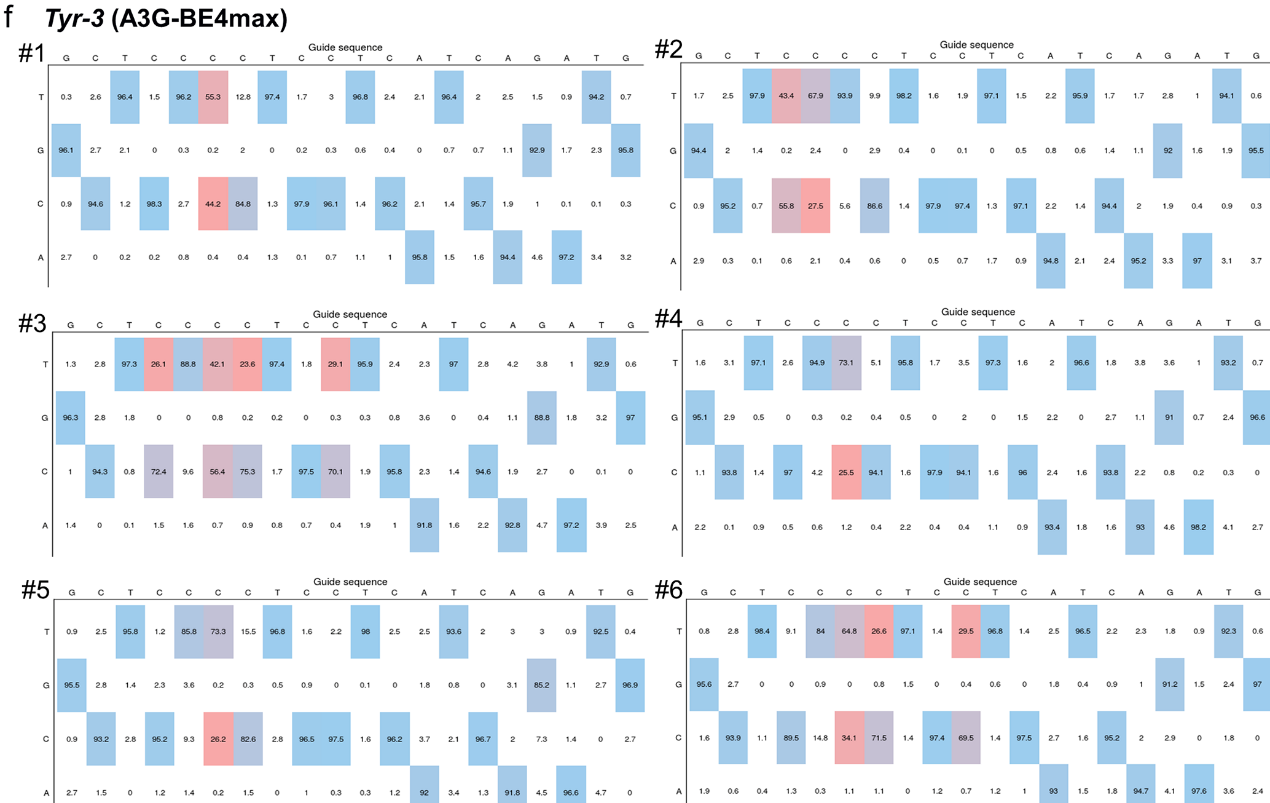

**Figure S4.** Base editing frequencies and product distribution at *Tyr-2* (**a**, **b**), *Fgf5* (**c**, **d**) and *Tyr-3* (**e**, **f**) sites with CCC context in rabbit blastocysts using BE4max and A3G-BE4max. The output editing tables in a colored heat map were produced by evaluating the Sanger sequencing results of each blastocyst by EditR.

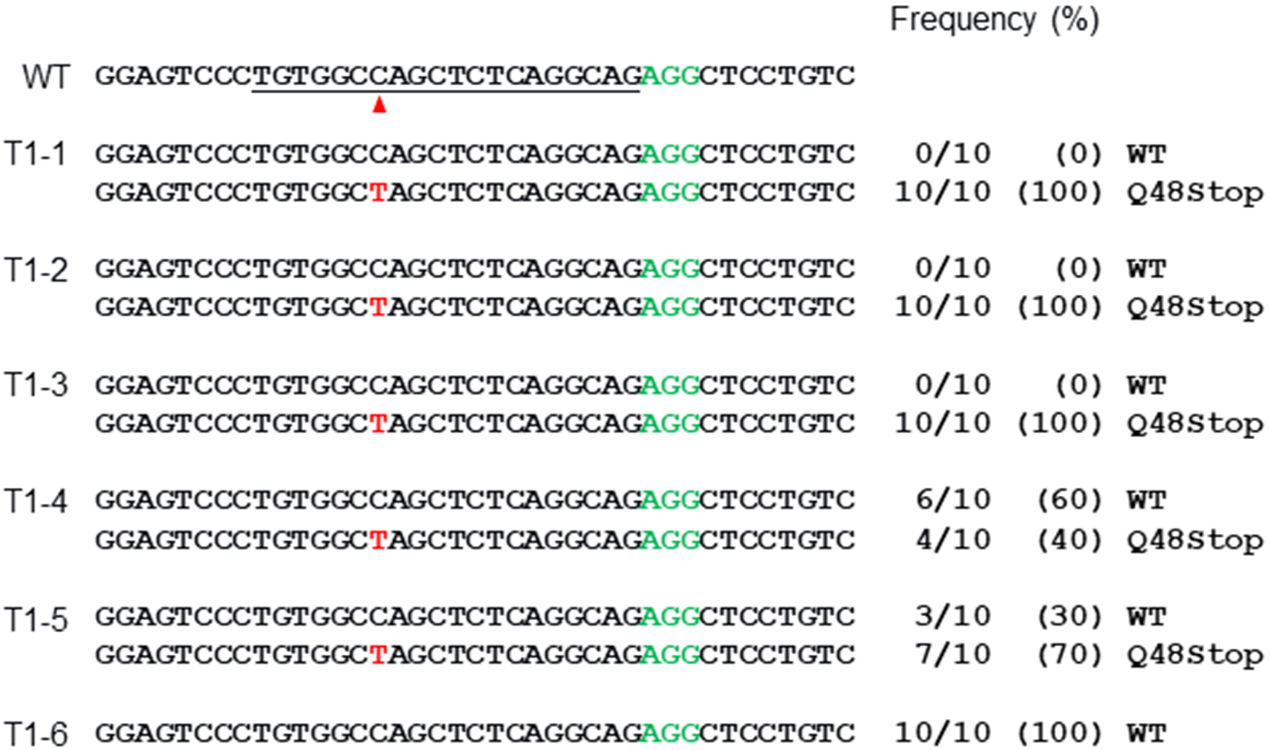

**Figure S5.** Alignments of mutant sequences of F0 rabbits from T-A cloning at *Tyr-1* using A3G-BE4max. The targeted sequence is underlined. The PAM site and base conversions are shown in green and red, respectively. The column on the right indicates frequencies of mutant alleles. WT, wild-type. T1-1 to T1-6, each individual.

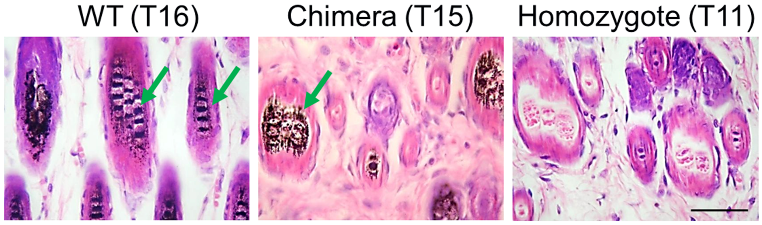

**Figure S6.** H&E staining of skin from WT (T16) and *Tyr-1* mutant (T11 and T15) rabbits. The green arrows highlight the melanin in the basal layer of the epidermis. Scale bars: 50 μm.

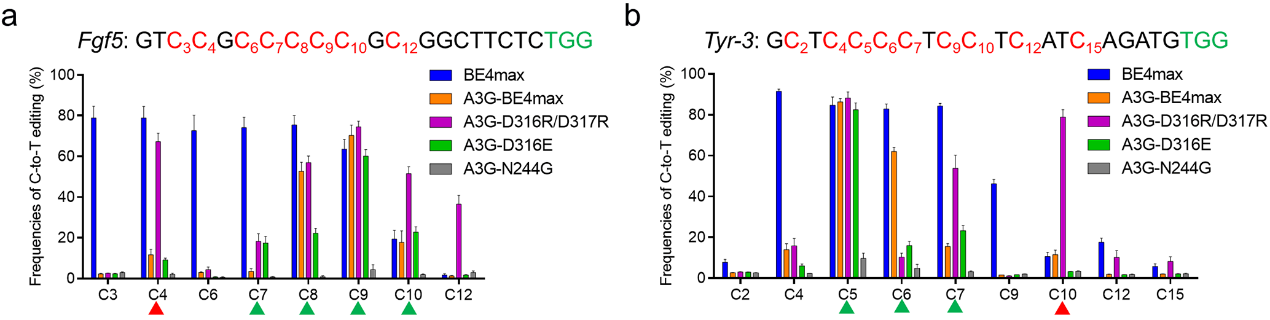

**Figure S7.** Summary of C-to-T base editing frequencies induced by BE4max and various A3G-BE4max variants on each cytosine at *Fgf5* (**a**) and *Tyr-3* (**b**). Targeted sequence (black), PAM region (green) and targeted Cs (red). Cytosines are counted with the base distal to the PAM setting as position 1. CC context (red triangle) and CCC context (green triangle)

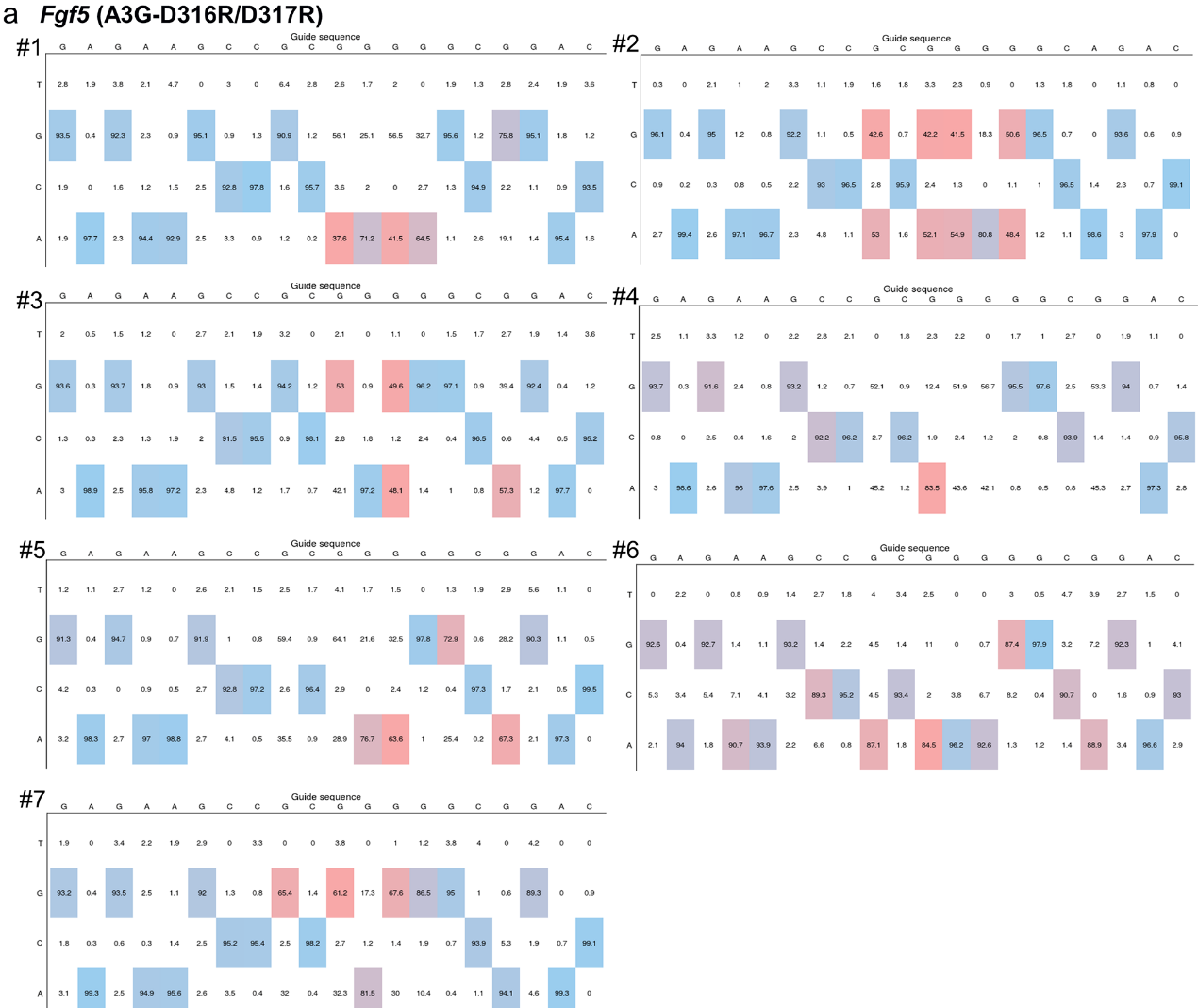

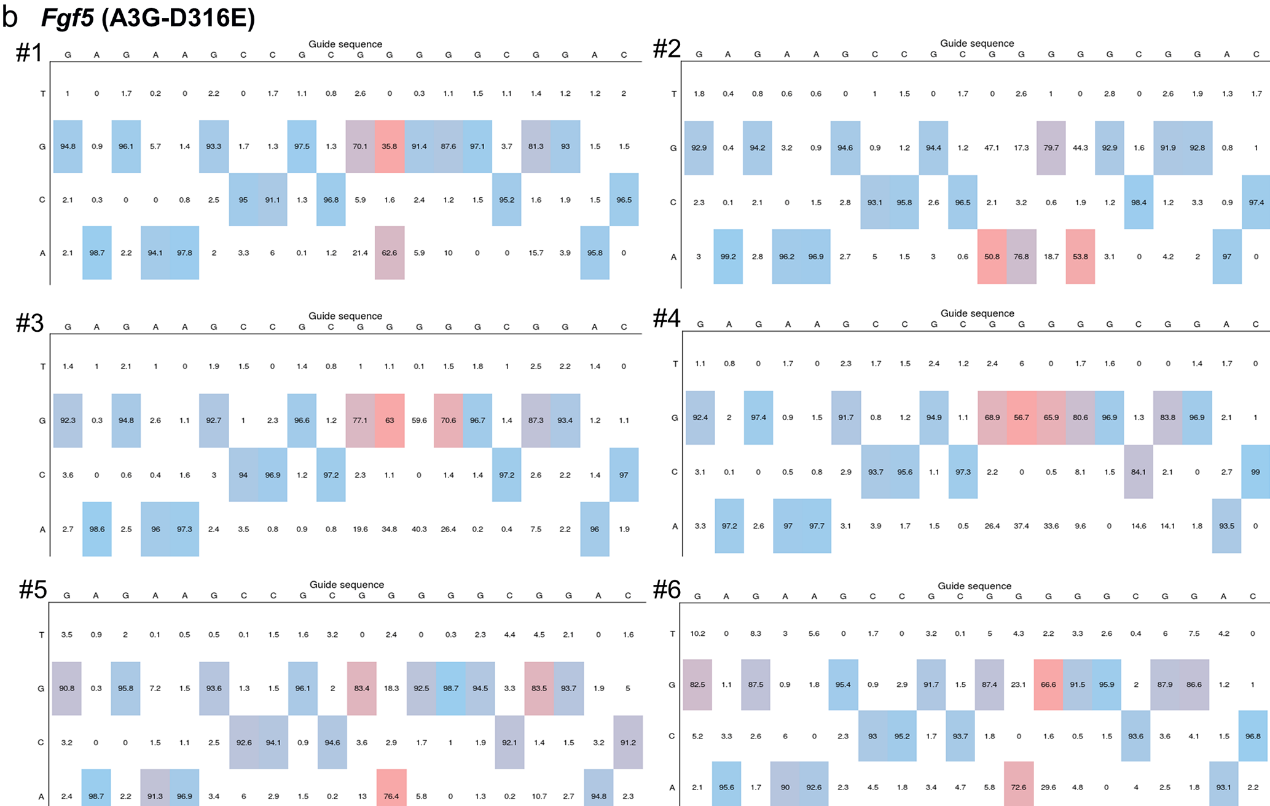

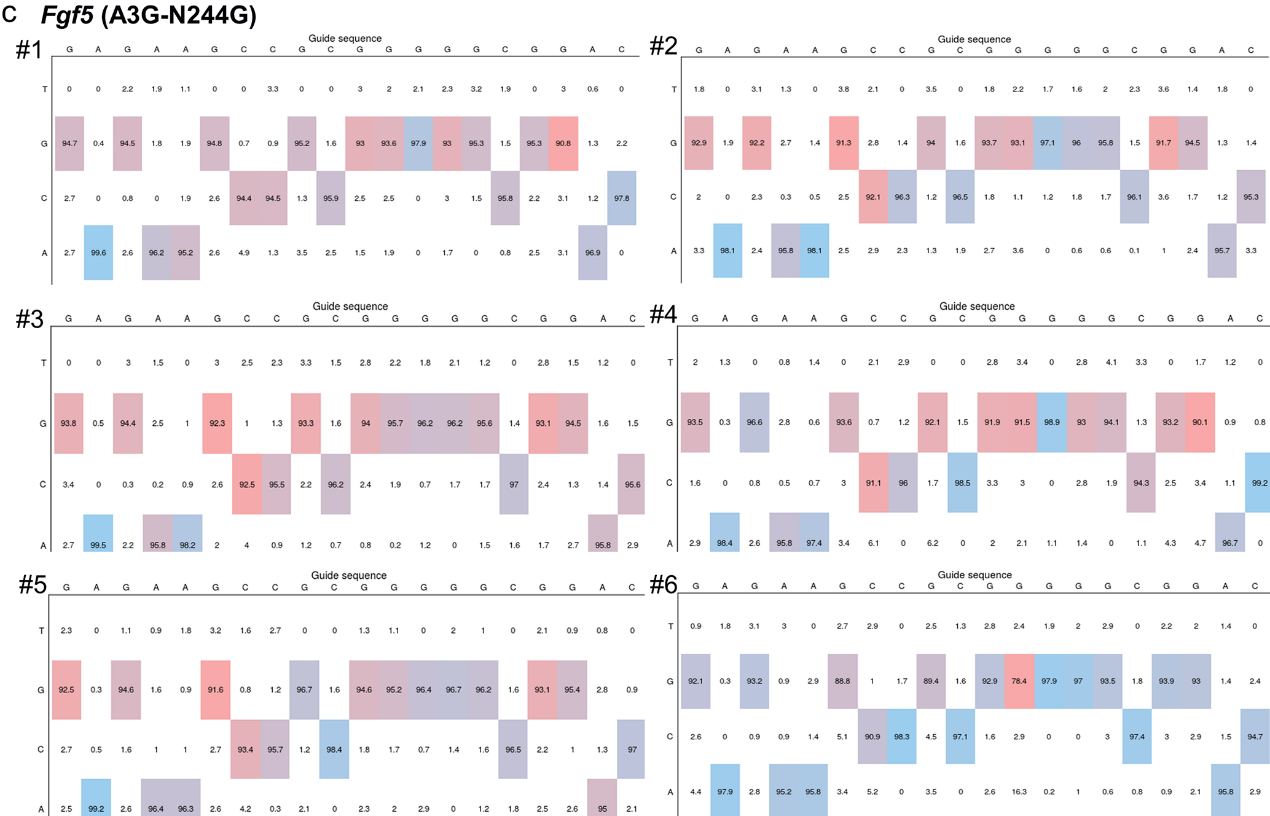

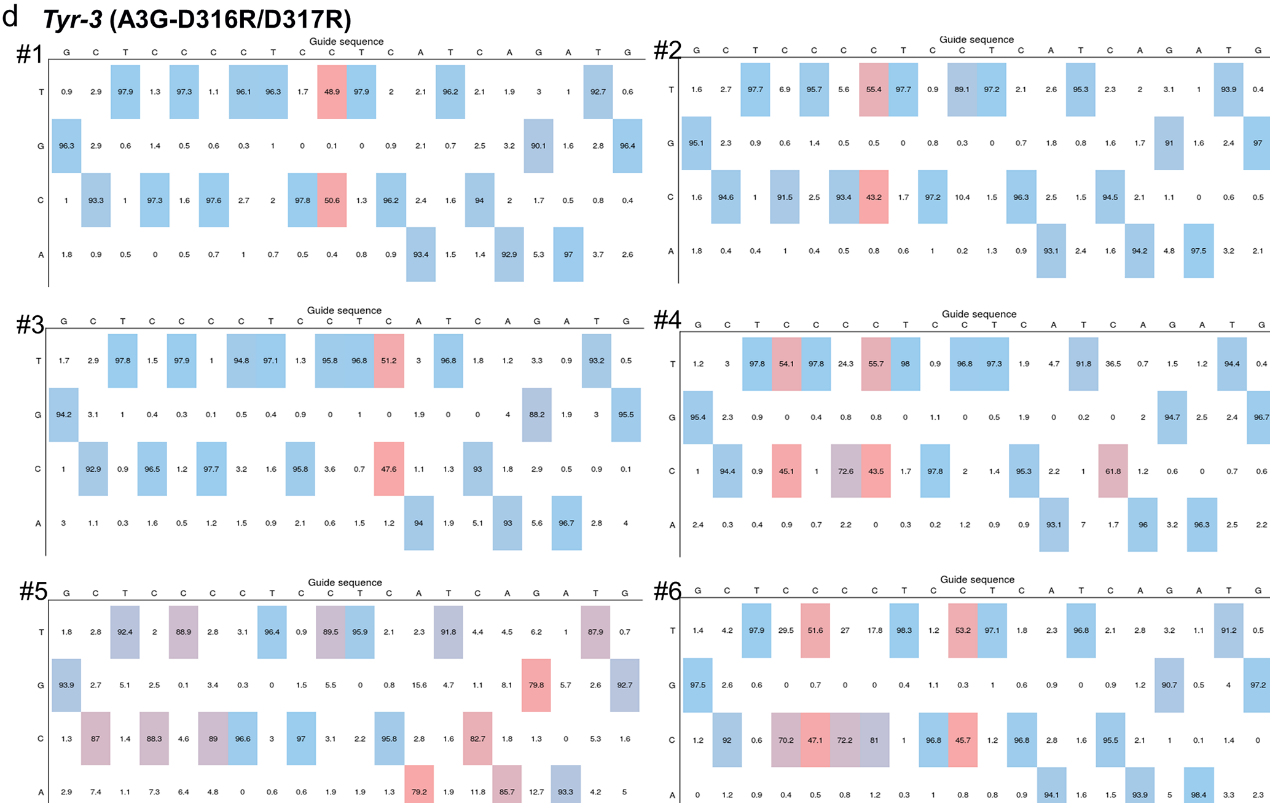

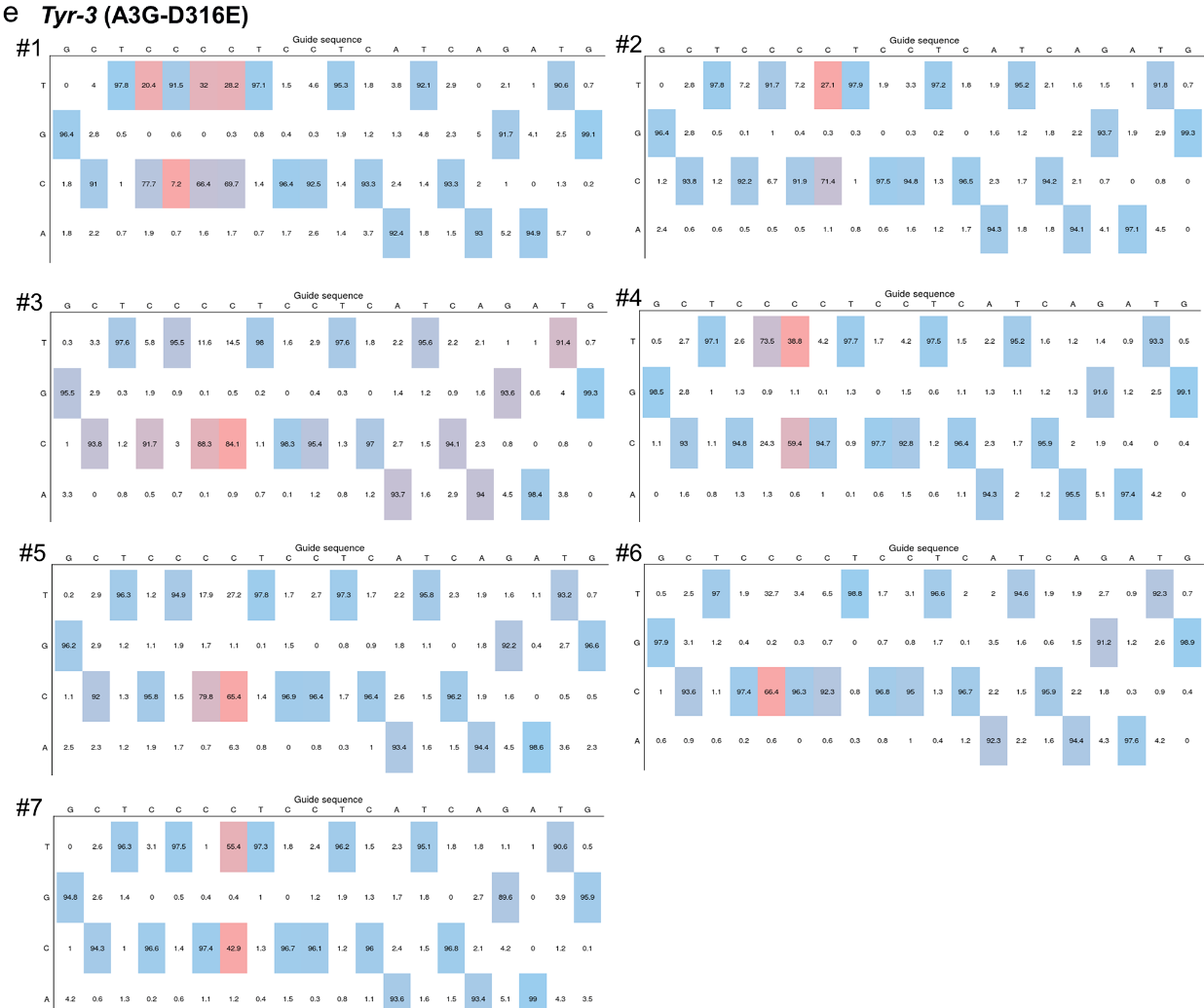

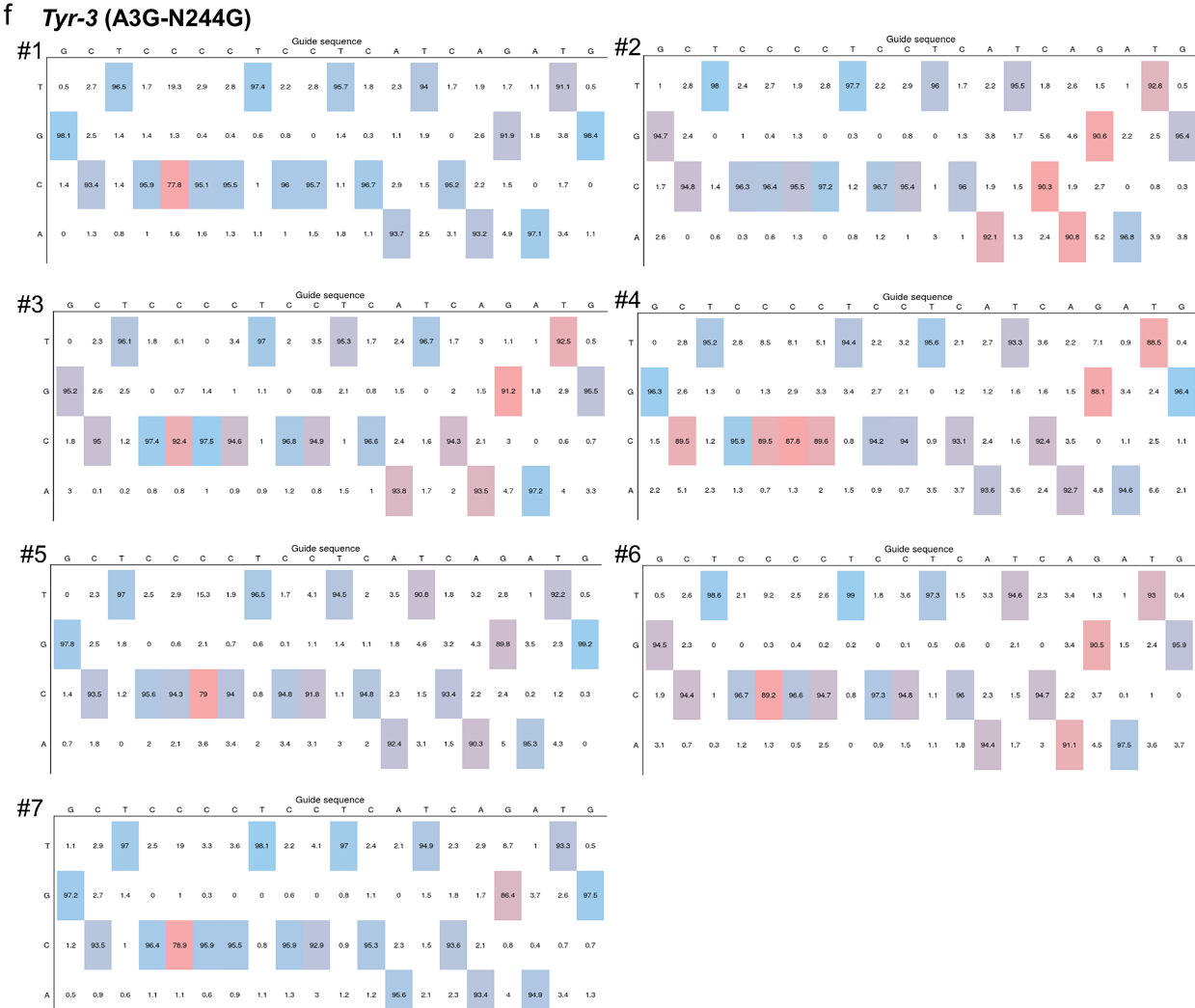

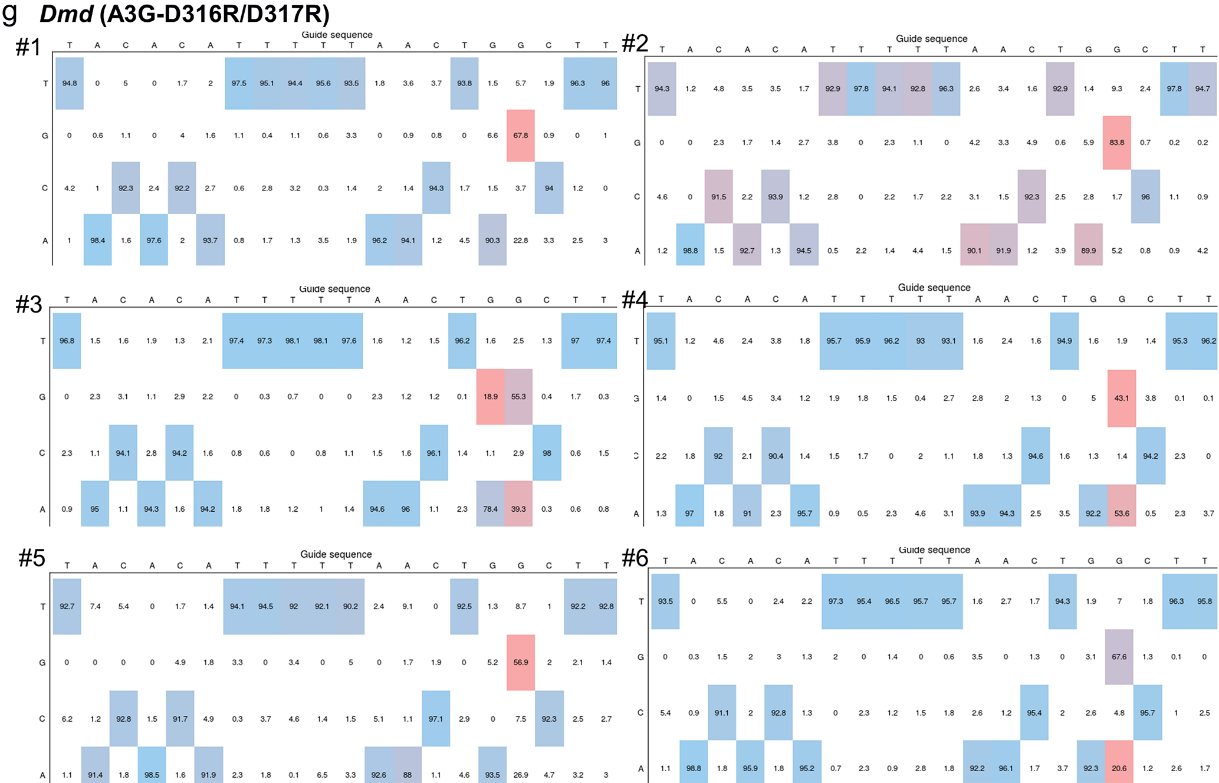

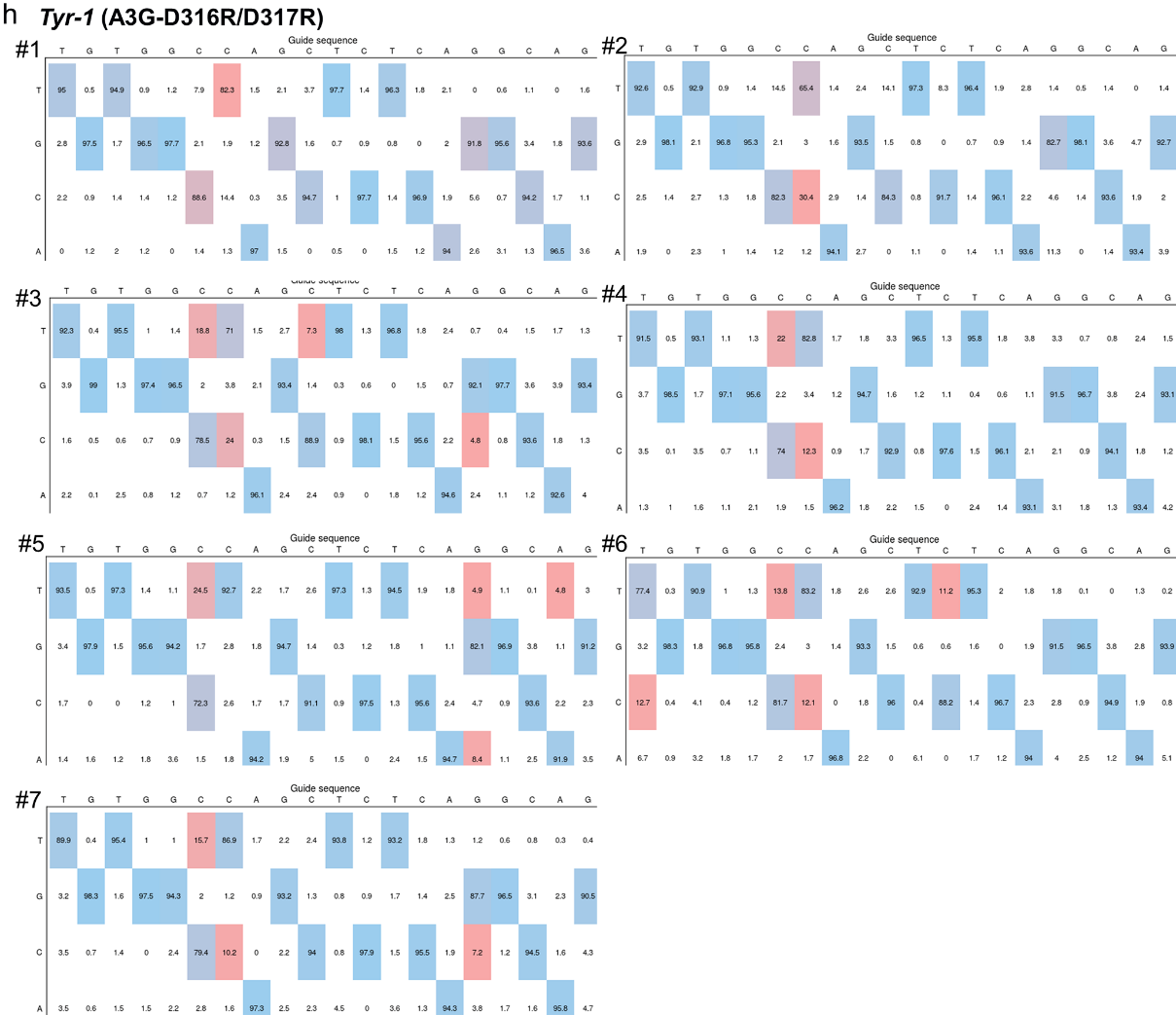

**Figure S8.** Base editing frequencies and product distribution at *Fgf5* (**a-c**), *Tyr-3* (**d**-**f**), *Dmd* (**g**) and *Tyr-1* (**h**) sites in rabbit blastocysts using various A3G-BE4max variants. The output editing tables in a colored heat map were produced by evaluating the Sanger sequencing results of each blastocyst by EditR.

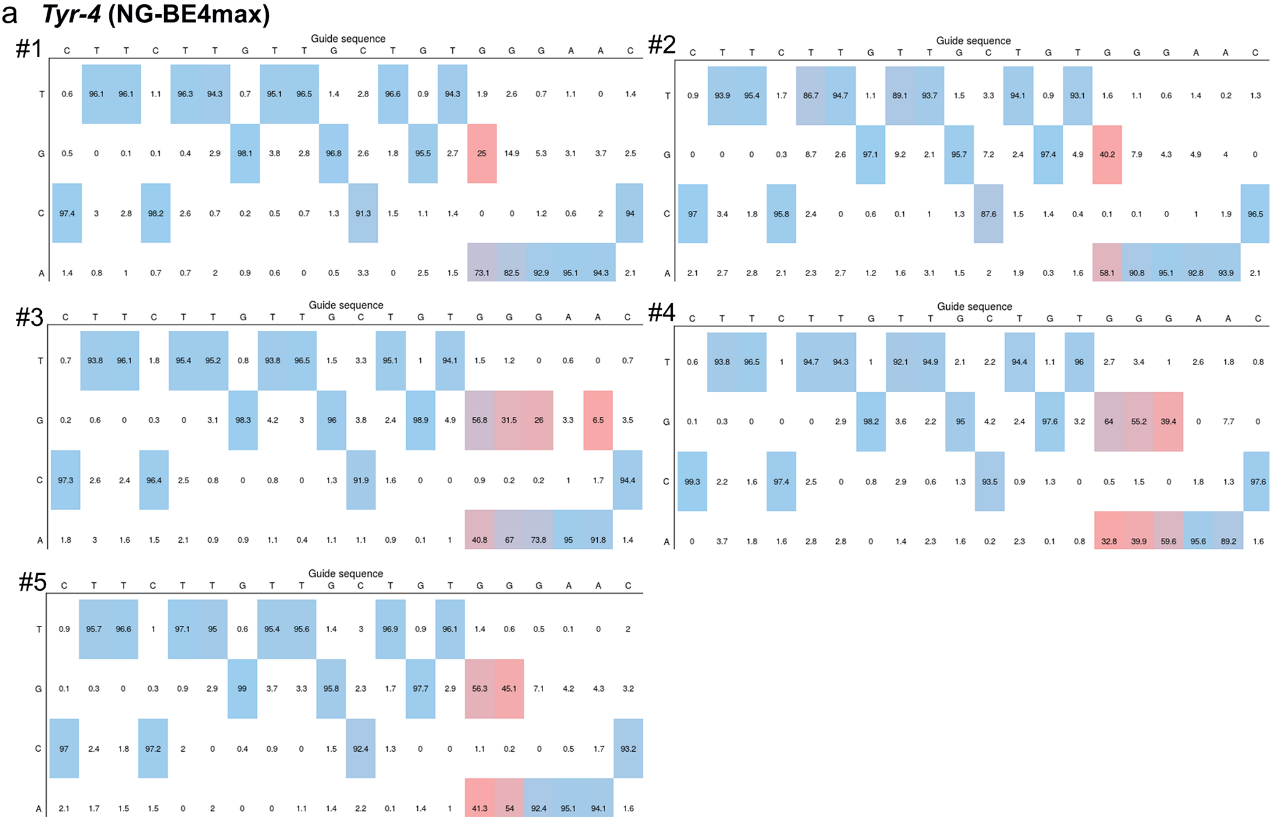

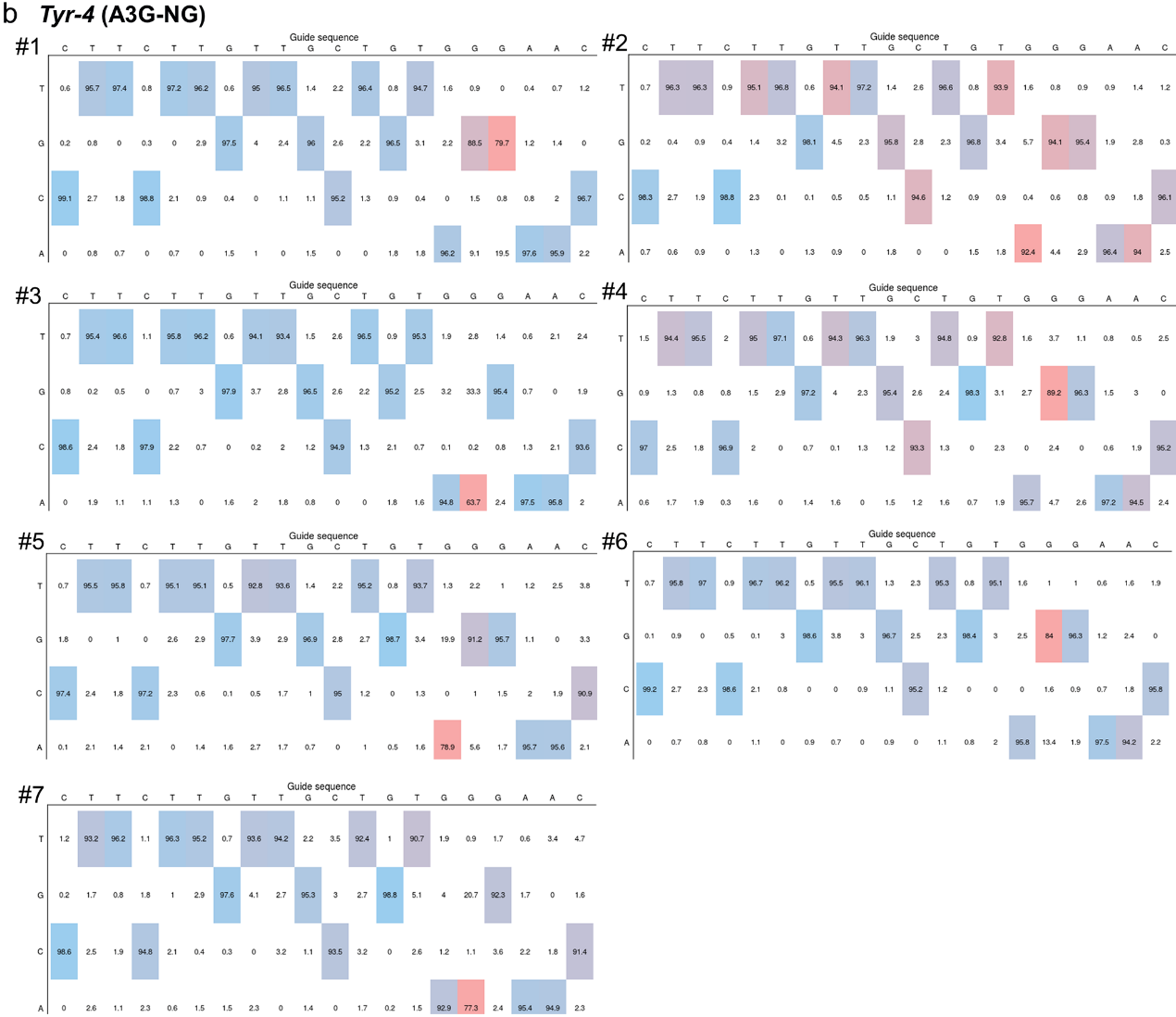

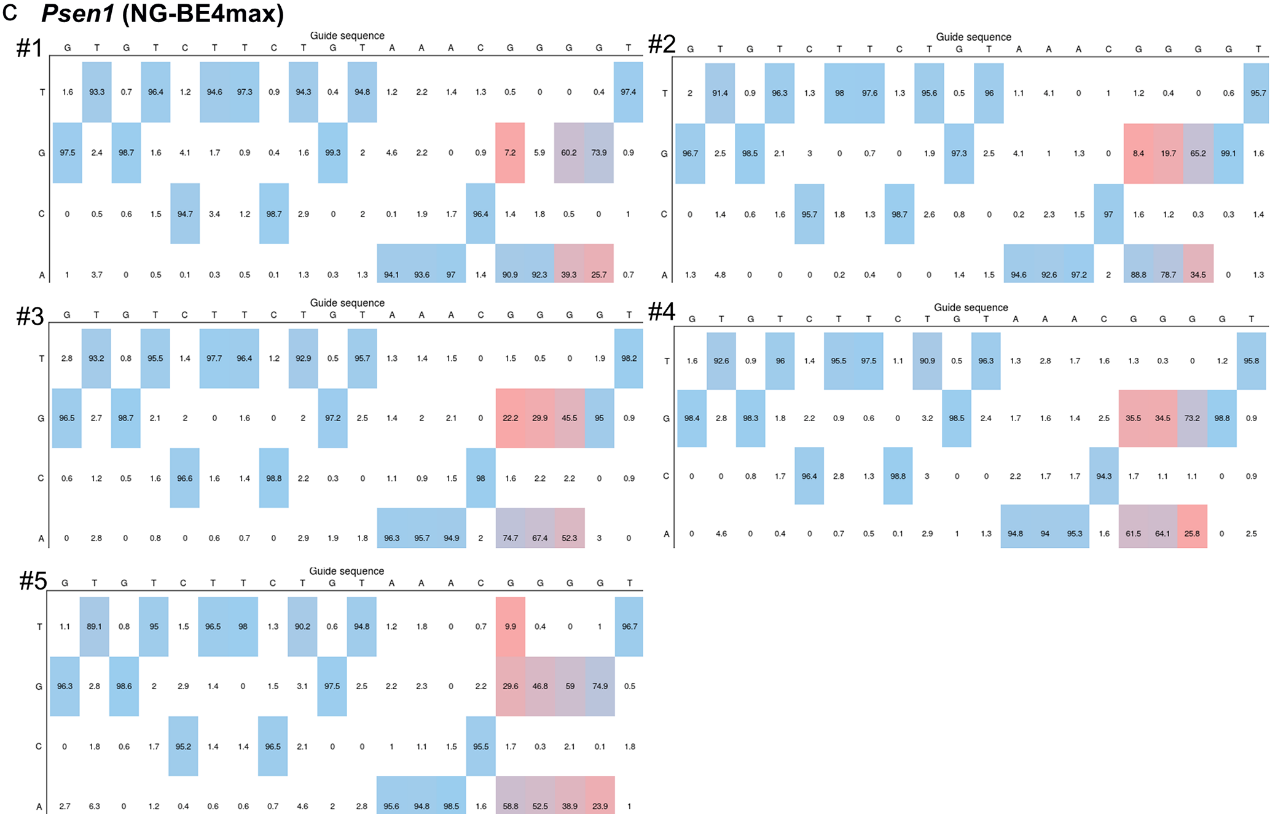

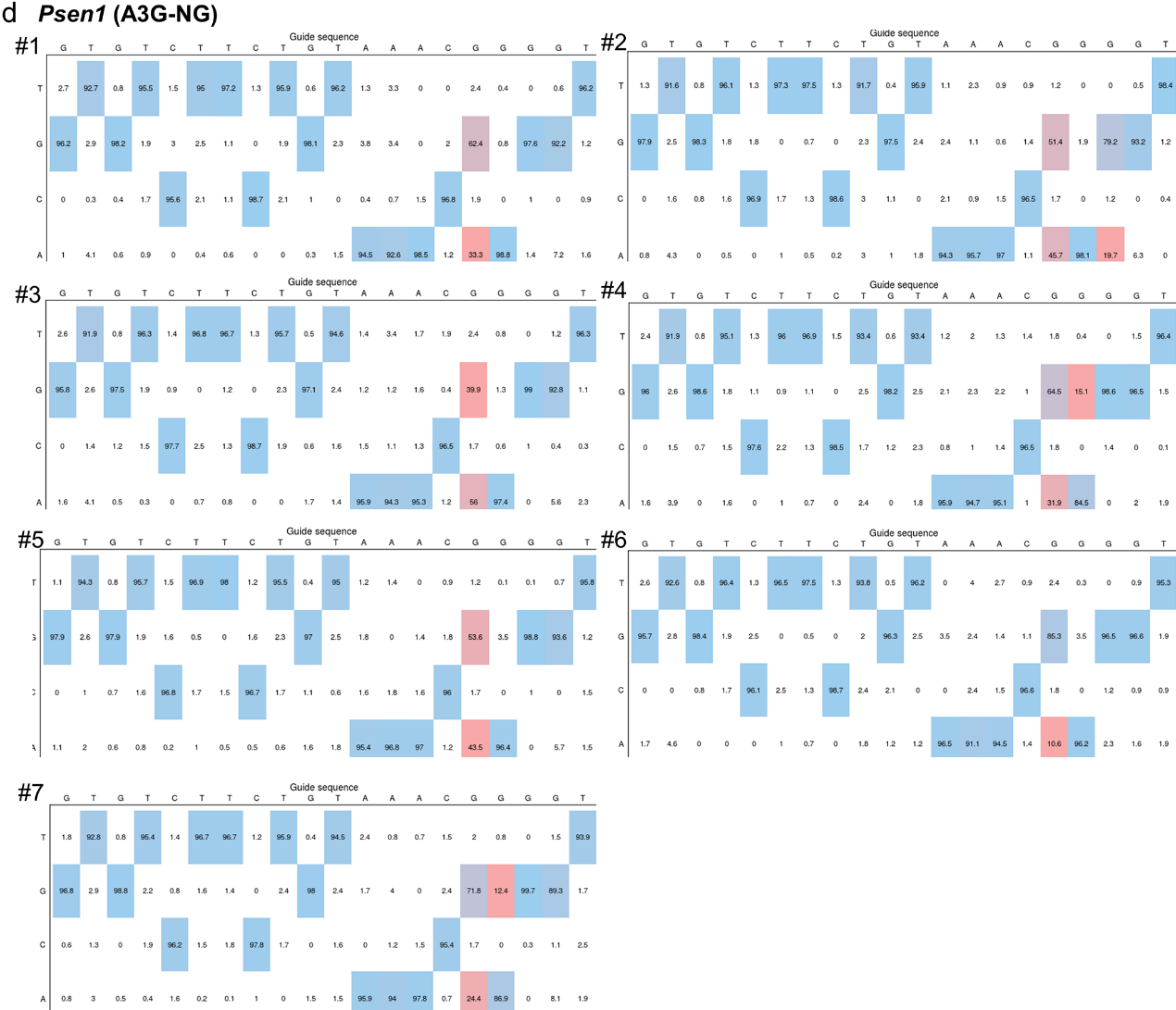

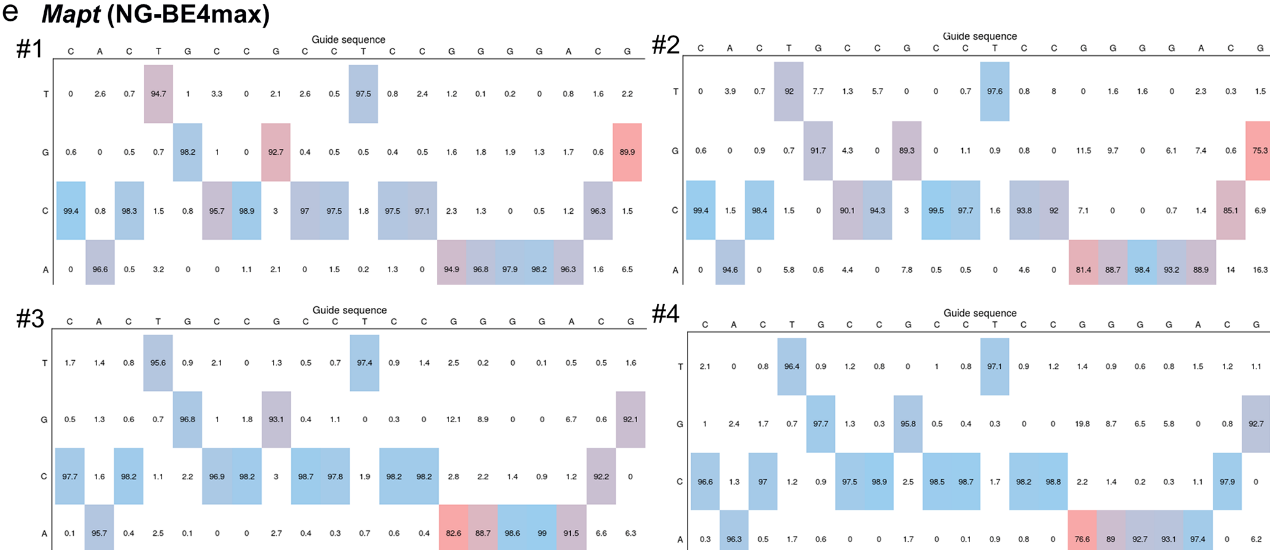

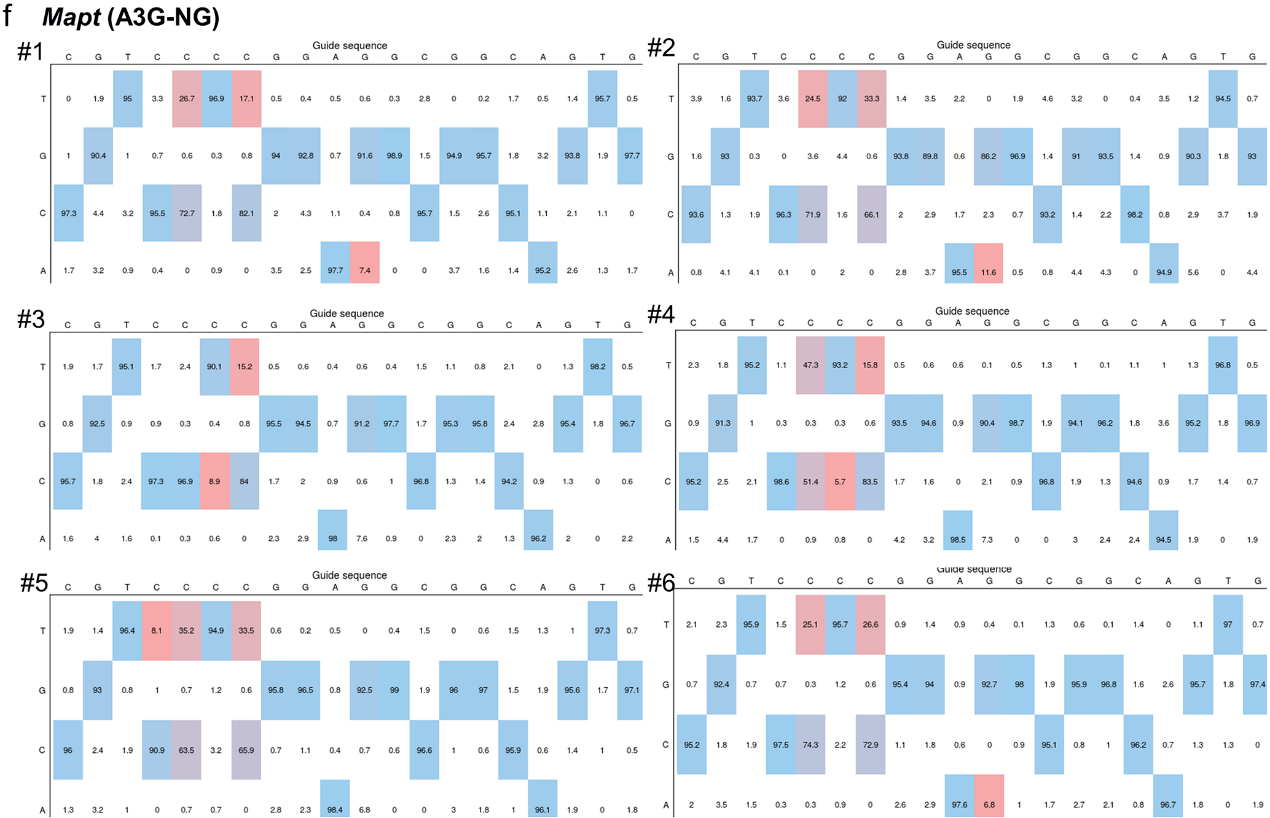

**Figure S9.** Base editing frequencies and product distribution at *Tyr-4* (**a**, **b**), *Psen1* (**c**, **d**) and *Mapt* (**e**, **f**) sites in rabbit blastocysts using NG-BE4max and A3G-NG. The output editing tables in a colored heat map were produced by evaluating the Sanger sequencing results of each blastocyst by EditR.

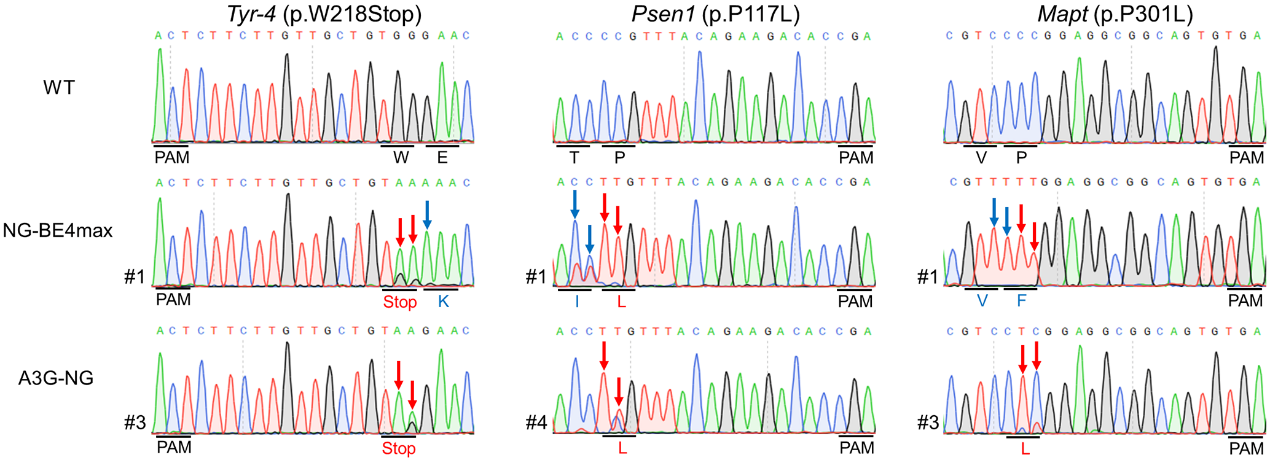

**Figure S10.** Representative sequencing chromatograms of edited rabbit blastocyst at three target sites using NG-BE4max and A3G-NG systems. Targeted mutations (red arrows) and bystander mutations (blue arrows). The relevant codon identities at the target site are presented under the DNA sequence.

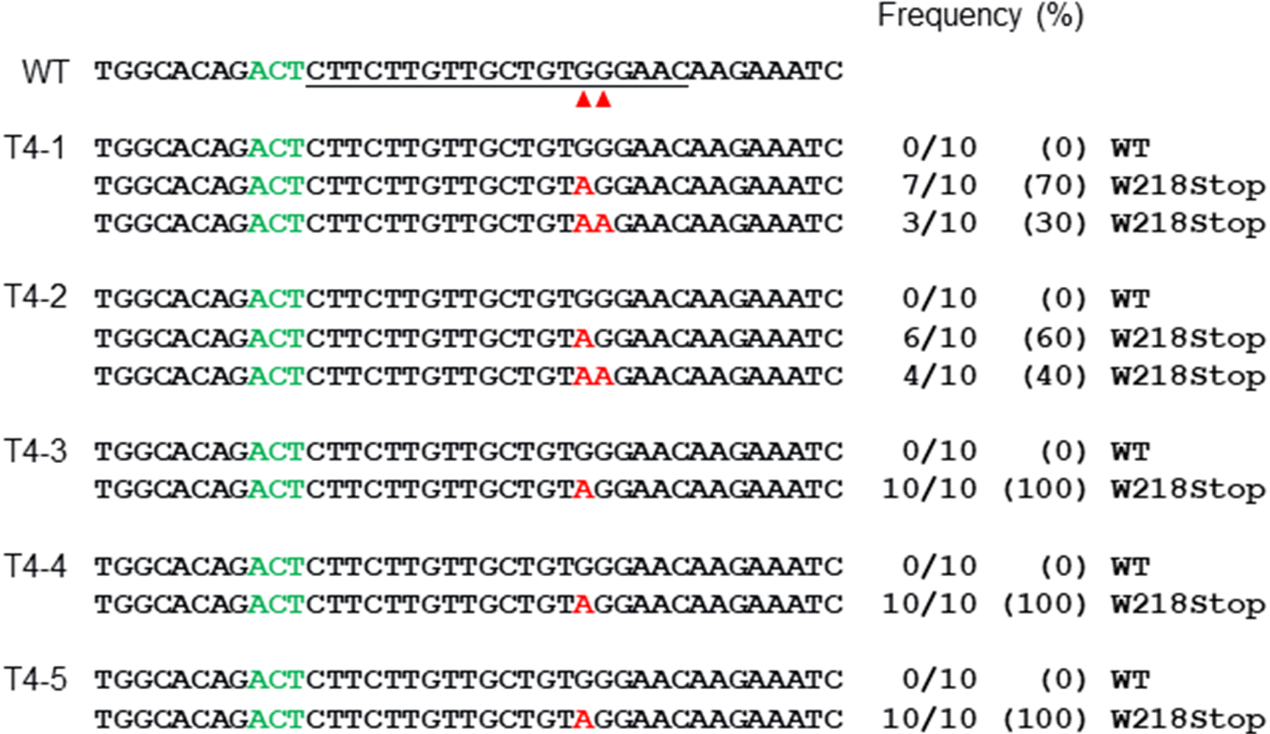

**Figure S11.** Alignments of mutant sequences of F0 rabbits from T-A cloning at *Tyr-4* using A3G-NG. The targeted sequence is underlined. The PAM site and base conversions are shown in green and red, respectively. The column on the right indicates frequencies of mutant alleles. WT, wild-type. T4-1 to T4-5, each individual.

**Table S1.** Target sites used in this study. Target sequence (black), PAM region (green).

| **PAM** | **Target site** | **Target site (5’-3’)** | **Pathogenic mutation** | **Human genetic disease** |
| --- | --- | --- | --- | --- |
| NGG | *Tia1* | CAGCCGCCTCCTGGCCAGAACGG | p.P362L | ALS |
| NGG | *Dmd* | AAGCCAGTTAAAAATGTGTAAGG | p.Q869Stop | DMD |
| NGG | *Tyr-1* | TGTGGCCAGCTCTCAGGCAGAGG | p.Q48Stop | OCA1 |
| NGG | *Tyr-2* | AGTCCCTGTGGCCAGCTCTCAGG | *NA* | *NA* |
| NGG | *Fgf5* | GTCCGCCCCCGCGGCTTCTCTGG | *NA* | *NA* |
| NGG | *Tyr-3* | GCTCCCCTCCTCATCAGATGTGG | *NA* | *NA* |
| NGT | *Tyr-4* | ACTCTTCTTGTTGCTGTGGGAAC | p.W218Stop | OCA1 |
| NGA | *Psen1* | ACCCCGTTTACAGAAGACACCGA | p.P117L | AD |
| NGA | *Mapt* | CGTCCCCGGAGGCGGCAGTGTGA | p.P301L | AD |

*NA*, not applicable.

**Table S2.** Summary of F0 rabbits generated in this study.

|  |  |  |  | **Mutant ratio (%)** | | |
| --- | --- | --- | --- | --- | --- | --- |
| **System** | **Target site** | **Embryos**  **transferred** | **No. of offspring** | **No. of mutants** | **No. of**  **homozygous mutants** | **No. of**  **bystander mutants** |
| A3G-BE4max | *Tyr-1* | 42 | 6 | 5(83) | 3(50) | 0(0) |
| A3G-NG | *Tyr-4* | 38 | 5 | 5(100) | 5(100) | 0(0) |

**Table S3.** Primers used for site-directed mutation in this study.

| **Plasmid template** | **Mutation** | **Primers (5’-3’)** |
| --- | --- | --- |
| A3G-BE4max | D316R/D317R (A3G-CTD) | F: ACCGCCCGGATCTACCGGCGGCAGGGCAGGTGTCAG  R: CTGACACCTGCCCTGCCGCCGGTAGATCCGGGCGGT |
|  | D316E  (A3G-CTD) | F: CGCCCGGATCTACGAGGATCAGGGCAGGTG  R: CTCGTAGATCCGGGCGGTGAAGATGCACAG |
|  | N244G  (A3G-CTD) | F: AGAGGCTTTCTGTGCGGCCAGGCCCCTCAC  R: GCCGCACAGAAAGCCTCTCCGCTGATTCAG |
| BE4max | L1111R (Cas9) | F: AGCAAAGAGTCTATCCGGCCCAAGAGGAAC  R: CGGATAGACTCTTTGCTGAAGCCGCCTGTC |
|  | D1135V (Cas9) | F: AGTACGGCGGCTTCGTGAGCCCCACCGTGG  R: CACGAAGCCGCCGTACTTCTTAGGGTCCCA |
|  | G1218R,  E1219F (Cas9) | F: GAATGCTGGCCTCTGCCAGATTCCTGCAGAAGGGAAACGA  R: TCGTTTCCCTTCTGCAGGAATCTGGCAGAGGCCAGCATTC |
|  | A1322R (Cas9) | F: AATCTGGGAGCCCCTCGGGCCTTCAAGTAC  R: CCGAGGGGCTCCCAGATTGGTCAGGGTAAA |
|  | R1335A, T1337R (Cas9) | F: CACCATCGACCGGAAGGTGTACCGGAGCACCAAAGAGGTG  R: CACCTCTTTGGTGCTCCGGTACACCTTCCGGTCGATGGTG |

**Table S4.** The two oligonucleotide strands used to construct the pUC57-sgRNA vectors.

| **Target site** | **Oligonucleotide 1** | **Oligonucleotide 2** |
| --- | --- | --- |
| *Tia1* | TAGGCAGCCGCCTCCTGGCCAGAA | AAACTTCTGGCCAGGAGGCGGCTG |
| *Dmd* | TAGGAAGCCAGTTAAAAATGTGTA | AAACTACACATTTTTAACTGGCTT |
| *Tyr-1* | TAGGTGTGGCCAGCTCTCAGGCAG | AAACCTGCCTGAGAGCTGGCCACA |
| *Tyr-2* | TAGGAGTCCCTGTGGCCAGCTCTC | AAACGAGAGCTGGCCACAGGGACT |
| *Fgf5* | TAGGTCCGCCCCCGCGGCTTCTC | AAACGAGAAGCCGCGGGGGCGGA |
| *Tyr-3* | TAGGCTCCCCTCCTCATCAGATG | AAACCATCTGATGAGGAGGGGAG |
| *Tyr-4* | TAGGTTCCCACAGCAACAAGAAG | AAACCTTCTTGTTGCTGTGGGAA |
| *Psen1* | TAGGACCCCGTTTACAGAAGACAC | AAACGTGTCTTCTGTAAACGGGGT |
| *Mapt* | TAGGCGTCCCCGGAGGCGGCAGTG | AAACCACTGCCGCCTCCGGGGACG |

**Table S5.** Primers used for genotyping in this study.

| **Target site** | **Primers** | **Sequence (5’-3’)** | **Product size (bp)** |
| --- | --- | --- | --- |
| *EMX1* | *EMX1*-F  *EMX1*-R | CAGCTCAGCCTGAGTGTTGA  CTCGTGGGTTTGTGGTTGC | 277 |
| *FANCF* | *FANCF*-F  *FANCF*-R | CATTGCAGAGAGGCGTATCA  GGGGTCCCAGGTGCTGAC | 182 |
| Site A | Site A-F  Site A-R | CAGCTCAGCCTGAGTGTTGA  CTCGTGGGTTTGTGGTTGC | 277 |
| *Tia1* | *Tia1*-F  *Tia1*-R | GGCATTACTGTTACGTTGGTATTT GGCAGACATCCAGCATCTT | 459 |
| *Dmd* | *Dmd*-F  *Dmd*-R | TCTTTCAGCCTGTGACTTCAG  GTGGCTTAGCTAAATCTGTAGGA | 421 |
| *Tyr-1* | *Tyr-1*-F  *Tyr-1*-R | ATCCGCTCAAGCAGGTATTG  GACATAGTCTGGGCTCGTAGTA | 487 |
| *Tyr-2* | *Tyr-2*-F  *Tyr-2*-R | ATCCGCTCAAGCAGGTATTG  GACATAGTCTGGGCTCGTAGTA | 487 |
| *Fgf5* | *Fgf5*-F  *Fgf5*-R | CTTCTTCAGCCACCTGATCTT  GGCGCAAGCAACTTACTTAAC | 342 |
| *Tyr-3* | *Tyr-3*-F  *Tyr-3*-R | GTGGTGGATGCAAGACTAGAA  AGCTGAAATTGGCAGCTTTG | 408 |
| *Tyr-4* | *Tyr-4*-F  *Tyr-4*-R | GCGACTCTTGGTGAGGAAA  AAAGATGCTGGGCTGAGTAG | 459 |
| *Psen1* | *Psen1*-F  *Psen1*-R | TTTACCTAGGGCTCTTGTGTTT  GTGTCTCAGGCTCACCTTATAG | 200 |
| *Mapt* | *Mapt*-F  *Mapt*-R | CCAGACTCTCCCAAGATTCTAATG  ACCCAGGCCGCTTTATTC | 468 |

**Supplementary sequence**

**Amino acid sequence of A3G-BE4max.**

Within the sequences below, NLS sequences are in purple, A3G-CTD sequences are in yellow (N244 (red), D316 and D317 (green)), the SpCas9 nickase sequence is in gray, the UGI sequences are in green.

MKRTADGSEFESPKKKRKVDPPTFTFNFNNEPWVRGRHETYLCYEVERMHNDTWVLLNQRRGFLCNQAPHKHGFLEGRHAELCFLDVIPFWKLDLDQDYRVTCFTSWSPCFSCAQEMAKFISKNKHVSLCIFTARIYDDQGRCQEGLRTLAEAGAKISIMTYSEFKHCWDTFVDHQGCPFQPWDGLDEHSQDLSGRLRAILQNQENSGGSSGGSSGSETPGTSESATPESSGGSSGGSDKKYSIGLAIGTNSVGWAVITDEYKVPSKKFKVLGNTDRHSIKKNLIGALLFDSGETAEATRLKRTARRRYTRRKNRICYLQEIFSNEMAKVDDSFFHRLEESFLVEEDKKHERHPIFGNIVDEVAYHEKYPTIYHLRKKLVDSTDKADLRLIYLALAHMIKFRGHFLIEGDLNPDNSDVDKLFIQLVQTYNQLFEENPINASGVDAKAILSARLSKSRRLENLIAQLPGEKKNGLFGNLIALSLGLTPNFKSNFDLAEDAKLQLSKDTYDDDLDNLLAQIGDQYADLFLAAKNLSDAILLSDILRVNTEITKAPLSASMIKRYDEHHQDLTLLKALVRQQLPEKYKEIFFDQSKNGYAGYIDGGASQEEFYKFIKPILEKMDGTEELLVKLNREDLLRKQRTFDNGSIPHQIHLGELHAILRRQEDFYPFLKDNREKIEKILTFRIPYYVGPLARGNSRFAWMTRKSEETITPWNFEEVVDKGASAQSFIERMTNFDKNLPNEKVLPKHSLLYEYFTVYNELTKVKYVTEGMRKPAFLSGEQKKAIVDLLFKTNRKVTVKQLKEDYFKKIECFDSVEISGVEDRFNASLGTYHDLLKIIKDKDFLDNEENEDILEDIVLTLTLFEDREMIEERLKTYAHLFDDKVMKQLKRRRYTGWGRLSRKLINGIRDKQSGKTILDFLKSDGFANRNFMQLIHDDSLTFKEDIQKAQVSGQGDSLHEHIANLAGSPAIKKGILQTVKVVDELVKVMGRHKPENIVIEMARENQTTQKGQKNSRERMKRIEEGIKELGSQILKEHPVENTQLQNEKLYLYYLQNGRDMYVDQELDINRLSDYDVDHIVPQSFLKDDSIDNKVLTRSDKNRGKSDNVPSEEVVKKMKNYWRQLLNAKLITQRKFDNLTKAERGGLSELDKAGFIKRQLVETRQITKHVAQILDSRMNTKYDENDKLIREVKVITLKSKLVSDFRKDFQFYKVREINNYHHAHDAYLNAVVGTALIKKYPKLESEFVYGDYKVYDVRKMIAKSEQEIGKATAKYFFYSNIMNFFKTEITLANGEIRKRPLIETNGETGEIVWDKGRDFATVRKVLSMPQVNIVKKTEVQTGGFSKESILPKRNSDKLIARKKDWDPKKYGGFDSPTVAYSVLVVAKVEKGKSKKLKSVKELLGITIMERSSFEKNPIDFLEAKGYKEVKKDLIIKLPKYSLFELENGRKRMLASAGELQKGNELALPSKYVNFLYLASHYEKLKGSPEDNEQKQLFVEQHKHYLDEIIEQISEFSKRVILADANLDKVLSAYNKHRDKPIREQAENIIHLFTLTNLGAPAAFKYFDTTIDRKRYTST

KEVLDATLIHQSITGLYETRIDLSQLGGDSGGSGGSGGSTNLSDIIEKETGKQLVIQESILMLPEEVEEVIGNKPESDILVHTAYDESTDENVMLLTSDAPEYKPWALVIQDSNGENKIKMLSGGSGGSGGSTNLSDIIEKETGKQLVIQESILMLPEEVEEVIGNKPESDILVHTAYDESTDENVMLLTSDAPEYKPWALVIQDSNGENKIKMLSGGSKRTADGSEFEPKKKRKV
